# Targeted high throughput mutagenesis of the human spliceosome reveals its *in vivo* operating principles

**DOI:** 10.1101/2022.11.13.516350

**Authors:** Irene Beusch, Beiduo Rao, Michael Studer, Tetiana Luhovska, Viktorija Šukytė, Susan Lei, Juan Oses-Prieto, Em SeGraves, Alma Burlingame, Stefanie Jonas, Hiten D. Madhani

## Abstract

The spliceosome is a staggeringly complex machine comprising, in humans, 5 snRNAs and >150 proteins. We scaled haploid CRISPR-Cas9 base editing to target the entire human spliceosome and interrogated the mutants using the U2 snRNP/SF3b inhibitor, pladienolide B. Hypersensitive substitutions define functional sites in the U1/U2-containing A-complex but also in components that act as late as the second chemical step after SF3b is dissociated. Viable resistance substitutions map not only to the pladienolide B binding site but also to the G-patch (ATPase activator) domain of SUGP1, which lacks orthologs in yeast. We used these mutants and biochemical approaches to identify the spliceosomal disassemblase DHX15/hPrp43 as the ATPase ligand for SUGP1. These and other data support a model in which SUGP1 promotes splicing fidelity by triggering early spliceosome disassembly in response to kinetic blocks. Our approach provides a template for the analysis of essential cellular machines in humans.

## INTRODUCTION

Pre-mRNA splicing is an essential step in eukaryotic gene expression. In addition to driving proteome diversity via alternative splicing (Blencowe, 2017), splicing impacts RNA stability, for example through the inclusion of exons with premature termination codons subject to nonsense-mediated decay (NMD), and plays critical roles in RNA export and translation efficiency (Le Hir et al., 2016). Splicing is also a major player in human disease: a large fraction of single nucleotide polymorphisms associated with human disease impact splicing (Li et al., 2016), and many human cancers harbor driver mutations in components of the spliceosome itself (Bejar, 2016; Yoshimi et al., 2019)

There are four intron sequences important for splicing: the 5’ splice site (SS), the branchpoint (BP), the polypyrimidine tract (PPT), and 3’ splice site (Fig. 1). In humans, there is large variability in these sequences, which can have an enormous impact on splicing efficiency and regulation, enabling regulation by RNA binding proteins (RBPs). Pre-mRNA splicing proceeds via two transesterification reactions which are catalyzed by the spliceosome. Compared to the simplicity of the chemical steps, the spliceosome is staggeringly complex (Figure 1 A,B). Components include five small nuclear ribonucleoproteins (snRNPs –U1, U2, U4/U6, and U5) and numerous proteins that assemble onto the intron substrate and undergo several large rearrangements to form a catalytically active complex in which a U6 snRNA acts as an RNA catalyst (Wilkinson et al., 2020). In *S. cerevisiae,* from which much of our understanding has been developed, splicing of a single intron requires eight ATP-dependent steps and about 90 proteins. Human spliceosomes appear to contain about 60 additional proteins (Wahl et al., 2009a).

**Figure 1.**
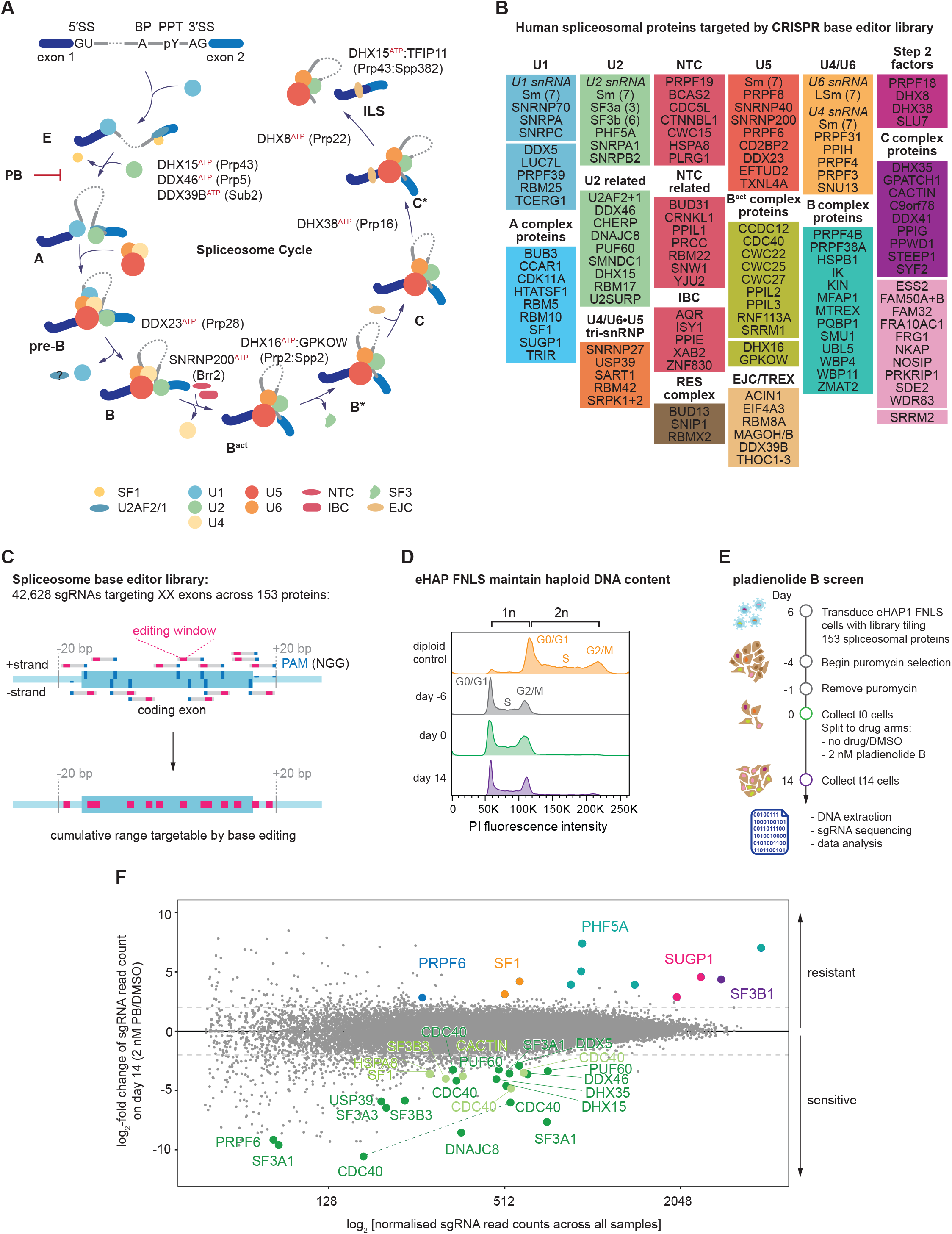
CRISPR-Cas9 base editing screen targeting the spliceosome reveals several mutants sensitive or resistant to the small molecule spliceosome inhibitor pladienolide B. (A) Schematic of intron sequences required for splicing and the spliceosome cycle across the major assembly stages. Depicted are involved snRNPs, important subcomplexes, proteins and helicases. Point of action of the small molecule spliceosome inhibitor pladienolide B (PB) is indicated. (B) List of spliceosomal genes targeted by the sgRNA library. (C) Schematic of tiling sgRNA library. Every available PAM sequence (denoted in dark blue) on both strands of the genome is targeted across all coding exons. CRISPR-Cas9 cytosine base editor is targeted to the genomic DNA through the sgRNA to promote C > T editing within each sgRNA’s editing window (marked in pink). This results in a theoretical cumulative range targetable by base editing. (D) eHAP FNLS cell line can be maintained in a haploid state. Shown is the flow cytometry analysis of DNA content via propidium iodide staining at various days corresponding to the control arm of our CRISPR-Cas9 base editing screen. (E) Schematic of the pooled screen. After transduction cells are given six days for mutagenesis by the CRISPR-Cas9 base editor. Cells are then split into treatment arms (DMSO vs. 2 nM PB) and collected after 14 days of culturing for DNA extraction and deep sequencing. (F) Results of screen. MA-plot comparing day 14 of cells grown in presence or absence of 2 nM PB. For orientation lines indicate a log_2_-fold enrichment or depletion of two. sgRNA with strong sensitivity to PB are emphasized in green (dark green: p-adj < 0.05, light green: p-adj ≥ 0.05 but highly depleted). sgRNAs resulting in PB resistance are colored by protein target. For clarity, only data points for sgRNAs that passed the confirmation assay are shown. Dashed line: sgRNA targeting the same position and predicted to result in identical mutational outcome. Dark green: statistically significantly depleted sgRNAs. Light green: strongly depleted sgRNAs.

Initial intron recognition involves base-pairing between the 5’ end of U1 snRNA and the 5’ SS and recognition of the branchpoint, PPT, and 3’ SS by sequence-specific RNA binding proteins: Splicing Factor 1 (SF1/Msl5) recognizes the branchpoint sequence while the two subunits of U2AF recognize the PPT and the conserved AG dinucleotide at the 3’ splice site. This forms an early, or E, complex that is the precursor to the A complex in which U2 snRNP binds to the intron, base-pairing with sequences around the branchpoint (the branchpoint sequence), replacing SF1 and U2AF. A triple snRNP, containing base-paired U4/U6 snRNAs together with the U5 snRNA, then joins the complex to form the pre-B complex which converts to the B complex by the departure of U1. Activation of the spliceosome occurs via ATP-dependent rearrangements that expels the U4 snRNP and several proteins (Wahl et al., 2009b), allowing the PRPF19/Prp19 complex (NTC) and NTC-related proteins (NTR) to join. This produces the B^act^ complex, in which U6 base-pairs with the 5’ splice site and U2 and U6 snRNAs base-pair to form the spliceosomal active site (Wilkinson et al., 2020). A component of U2 snRNP, the SF3 complex, which sequesters the U2-branchpoint helix away from the 5’ splice site, is then removed, allowing the U2-branchpoint helix to dock with the catalytic core. Association of additional proteins allow the chemical steps to proceed in the B* and C* catalytic complexes (Wilkinson et al., 2020). The ATPases DHX38/Prp16 and DHX8/Prp22 respectively remodel the active site after each chemical step (Wahl et al., 2009b). Following mRNA release, the helicase DHX15/Prp43 disassembles the spliceosome (Martin et al., 2002; Tsai et al., 2005). Like other DEAH-box helicases, DHX15/Prp43 is activated by a cognate G-patch protein, TFIP11/Spp382/Ntr1 (Tanaka et al., 2007).

Given the high variability in splicing signal sequences in humans, how the spliceosome distinguishes between cognate and noncognate sequences remains to be understood. A longstanding hypothesis suggests that the dynamic and complex nature of the spliceosome promotes the fidelity of splicing through kinetic proofreading while also permitting substrate flexibility and regulation. Evidence supporting this model came from a genetic screen in *S. cerevisiae* in which missense mutations in the ATPase Prp16 were identified as suppressors of a mutation in the branchpoint adenosine sequence (Burgess and Guthrie, 1993). Subsequent *in vitro* studies demonstrated that mutant pre-mRNA substrates that assemble into spliceosomes, but are kinetically slow at either chemical step, trigger spliceosome disassembly prior to completion of the reaction, a process termed “discard” (Koodathingal et al., 2010; Koodathingal and Staley, 2013; Mayas et al., 2010; Mayas et al., 2006; Semlow and Staley, 2012). Failure to perform catalysis prior to ATP hydrolysis by Prp16 (step 1) or Prp22 (step 2) produces a spliceosome that can be disassembled by Prp43. These ATPases have been proposed to act as molecular timers for productive movement through the splicing pathway (Koodathingal and Staley, 2013). The yeast studies used mutant pre-mRNA substrates because their signals are always very close to the optimal consensus (Irimia and Roy, 2008). Whether there are analogous or additional fidelity mechanisms that operate in animal cells is unknown.

The ability to perform forward genetic screens in haploid *S. cerevisiae* was critical for the studies on spliceosome fidelity outlined above as well as numerous other foundational studies of splicing. To adapt these methods to human cells, we describe here a strategy to mutagenize the spliceosome in fully haploid human cells by developing and deploying a CRISPR-Cas9 base editor sgRNA library that targets the entire human spliceosome. After mutagenesis, we interrogated the spliceosome using the potent inhibitor pladienolide B (PB), which targets U2 snRNP by binding to a pocket between the SF3B1 and PHF5A subunits of the SF3b complex, preventing stabilization of the U2-branchpoint RNA duplex (Cretu et al., 2018b; Cretu et al., 2021; Gamboa Lopez et al., 2021; Teng et al., 2017b; Wu et al., 2018). Validation and genomic sequencing revealed resistance mutations in SF3B1 and PHF5A in residues adjacent to the compound binding pocket. We mapped hypersensitive mutants to U2 snRNP components, but also to factors that act as late as the second chemical step, after SF3b has dissociated. Strikingly, we obtained resistance mutants in SUGP1, a spliceosomal G-patch protein of unknown function that lacks orthologs in yeast and is also a newly proposed tumor suppressor whose loss underpins the splicing changes induced by cancer-associated SF3B1 mutations (Alsafadi et al., 2021; Liu et al., 2020; Zhang et al., 2019). Our resistance mutations in SUGP1 map in or adjacent to its G-patch motif and modulate splicing changes triggered by PB. We describe biochemical experiments that reveal the spliceosomal disassembly ATPase DHX15/hPrp43 to be the biologically relevant direct target of the SUGP1 G-patch domain. We propose a unified model in which SUGP1/DHX15-mediated disassembly of kinetically-slowed early splicing complexes explains compound resistance as well as oncogenic aberrant splicing events resulting from SF3B1 and SUGP1 mutations. More broadly, our results demonstrate the feasibility and utility of the programmed generation of informative viable haploid alleles targeting a complex essential gene expression machine in human cells.

## RESULTS

### Large-scale mutagenesis of the human spliceosome

A major impediment to the study of the human spliceosome *in vivo* has been the inability to program point mutations in endogenous genes on a large scale. CRISPR-Cas9 technology now provides such opportunities (Anzalone et al., 2020). Due to its scalability and ability to introduce point mutations, we chose CRISPR-Cas9 base editing for a forward genetic screen of the spliceosome (Figure S1B). We first generated a monoclonal stable cell line expressing FNLS (Zafra et al., 2018a), a cytosine base editor, in an eHAP (Essletzbichler et al., 2014) haploid cell background (hereafter: eHAP FNLS). During clonal cell line generation, we maintained eHAP FNLS cells as haploid so that we could subsequently assign genotype-phenotype relationships (Figure 1D). We assessed editing efficiency on a set of standard targets used previously (Zafra et al., 2018b) (Figure S1C,D). eHAP FNLS cells demonstrated efficient editing (up to >90%) at expected positions within the editing window, which spans positions 3-8 [with position 21-23 being the protospacer adjacent motif (PAM)], and induced transversions (C > R editing) at high frequencies (>25%) in some cases. Transversion editing has been described previously; its extent is cell line-dependent (Sánchez-Rivera et al., 2022).

We designed a single guide RNA (sgRNA) library targeting a hand-curated list of 153 human spliceosomal proteins which engage at various steps of the splicing cycle (Figure 1A), and are reproducibly detected through mass spectrometry (MS), interaction studies, and/or visualized in structural biology studies (see Figure 1B, Table S1)(Sales-Lee et al., 2021). Given that base editing outcomes are not fully predictable, to maximize mutagenesis we targeted every available NGG PAM sequence across all annotated exons plus 20 bp flanking intronic sequence (Figure 1C). Our library includes 42,618 sgRNAs targeting 42,650 sites, including 8,426 sgRNAs that target genomic sites but are predicted to be non-editing with FNLS, and an additional 1,000 guides that do not target genomic sites (non-targeting sgRNAs) (Doench et al., 2016). The library can in principle mutagenize up to 30% of spliceosomal protein coding sequences (Figure S1E) and edits are predicted to result in in missense mutations in >50% of cases with an additional 20% of edits predicted to impact protein sequence (Figure S1F, and Methods for details on mutation outcome prediction).

We cloned this library into lentiviral vectors that express the sgRNA and associate each to a unique barcode and produced virus for transduction (Figure S1A) (Boettcher et al., 2019). We transduced the library into eHAP FNLS cells, incubated them for 6 days to allow for editing and selection of transduced cells, and then split the selected pool into treatment arms (control/DMSO vs. 2 nM PB, which approximates its EC_50_). We cultured cells for two weeks while maintaining a representation of 500 cells per sgRNA. On days 0, 8 and 14 we isolated genomic DNA and amplified and sequenced the sgRNA inserts; we also did this for the input plasmid library (Figure 1E). Using the sgRNA-linked barcode, we randomly assigned sgRNAs to two sample populations and depletion vs. enrichment of sgRNAs was analysed for both time and treatment using DESeq2 (Love et al., 2014).

We then compared the abundances of guide sequences in the population that differed in their predicted consequences versus the plasmid input control 14 days after transduction. Given that most spliceosomal proteins are essential for cell survival, we anticipated that a subset of induced mutations would be lethal or result in reduced viability, and that their sgRNAs would therefore be depleted over time. As expected, sgRNAs predicted to promote mutations with more severe consequences such as splice site mutations or creation of stop codons displayed the strongest depletion as a class, consistent with efficient and precise editing at many of those sites (Figure S1G). Conversely, guides predicted to be non-editing or to produce silent mutations were generally not depleted

By comparing guide sequence abundances for day 14 for 2 nM PB versus the matched DMSO control, we observed that several guides were enriched or depleted upon compound treatment of the population (Figure 1F). For validation, we selected the sgRNAs showing statistically significant enrichment or depletion (LFC > |2|, padj < 0.05) between the 2 nM PB sample and its matched control sample on day 14. To this list we added a subset of sgRNAs with high average enrichment/depletion but did not pass statistical significance (see Methods for details). To enrich for PB-specific phenotypes, we required that those guides depleted after PB treatment did not show depletion between t0 and t14 in untreated cells. This procedure yielded 19 candidate-enriched sgRNAs and 26 candidate-depleted sgRNAs. These sgRNAs and three non-targeting sgRNAs were then subjected to an arrayed dual-color competition assay in which cells transduced with virus encoding a non-targeting sgRNA or transduced with a candidate-depleted or -enriched sgRNA were marked with distinct fluorescent proteins, respectively (Figure 2A). The assay confirmed the response to PB treatment for 23/26 of the candidate-depleted sgRNAs and 11/19 of the candidate-enriched sgRNAs. Except for a sgRNA targeting *SF3B1* and another targeting *PRPF6,* only sgRNAs found to be statistically significantly enriched validated in the confirmation assay, supporting the utility of the statistical approach used (Figure 2B, S2A).

**Figure 2.**
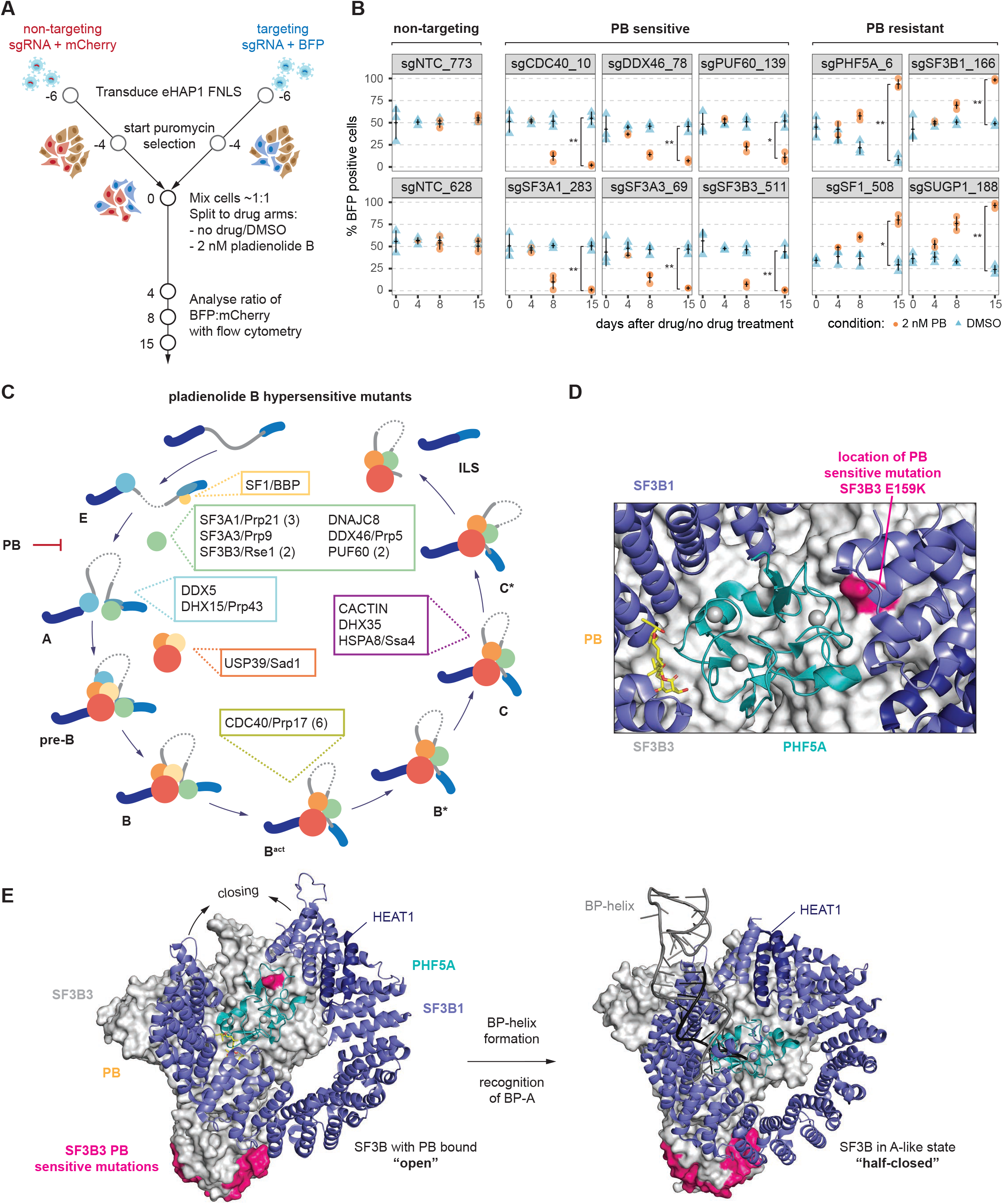
Pladienolide B sensitive mutations occur predominantly in early spliceosomal complexes. (A) Schematic of the arrayed confirmation assay. Individual sgRNA are transduced, marked with BFP, and cells are mixed 1:1 with non-targeting sgRNA carrying cells, marked by mCherry. Cell populations are left to compete during 15 days of growth in either DMSO or 2 nM PB treatment. The ratio of BFP:mCherry is measured with flow cytometry. (B) Individual sgRNAs and their performance in the confirmation assay. sgRNAs are grouped by category they were found in in the primary screen. Measurements are from three independent transductions (n=3). *; **; ***: Student’s t-Test (paired) *P* value < 0.05, 0.01, 0.001, respectively. (C) Assignment of proteins targeted by sgRNAs conferring hypersensitivity to PB to the spliceosome cycle. Factors are indicated at their respective first step of action. *Saccharomyces cerevisiae (Sc.)* names are given where applicable. If multiple sgRNAs are found for a protein, this is indicated with a number in parenthesis. (D) Close-up of location of PB-sensitive SF3A1 G159K mutation plotted on the structure of SF3b bound to PB. It lies at the interface of SF3B3 (green), SF3B1 (violet) and PHF5A (teal). (PDB: 6EN4) (E) Comparison of SF3B3, SF3B1 and PHF5A in the structure of SF3b bound to PB and the A-like complex. SF3B1 undergoes a large conformational rearrangement from the open to the half-closed state. For easier tracking of the conformational change, HEAT repeat 1 (HEAT1) is indicated. Locations of PB-sensitive mutations are marked in magenta. (PDB: 6EN4, 7Q4O)

### Pladienolide B hypersensitive mutations identify functional spliceosomal residues

Given the notable number of sgRNAs depleted upon compound treatment, we determined the genomic consequences of base editing. We transduced eHAP FNLS with lentivirus carrying individual sgRNAs, grew cells for six days, isolated genomic DNA, amplified the edited locus and subjected the amplicons to deep sequencing. This approach revealed the consequences of base editing at the amino acid level (Table S2).

Guides presumably must edit efficiently to produce a depletion phenotype. Amplicon sequence confirmed this expectation: all sgRNAs displayed high/substantial rates of editing (median = 57.7% for C > T within positions 3 to 8, Figure S2B) leading to amino acid changes that become depleted in the presence of PB (with a median of 82% of the sequences carrying an amino acid change, Table S2). Again, we observed not only C > T editing but also C > R editing, with predicted mutations matching for 19/23 sgRNAs. Thirteen distinct guides programmed mutations in early-acting spliceosomal factors, including SF1, SF3, and DDX46/hPrp5 as well as the U2AF-associated DEAD box protein DDX5/UAP56 (Figure 2C). Unexpectedly, we also identified a mutation in the tri-snRNP-specific protein USP39/hSad1 and five mutations in the second-step factor CDC40/hPrp17, which is first found in the B^act^ complex (Haselbach et al., 2018). Finally, we found PB-sensitive mutations in factors that join catalytically active complexes and act at the second chemical step of splicing (DHX35/hPrp16 and CACTIN), a point in the spliceosome cycle after which SF3b has been dissociated. As a first step in understanding how these changes impact the spliceosome, below we briefly place some of these mutations into the context of existing spliceosome structures, focusing on the SF3 complex.

PB sterically blocks binding of the U2-intron branchpoint duplex to its pocket in SF3b (Cretu et al., 2021), which is necessary for spliceosome assembly to proceed beyond the A complex. Six of our PB-sensitive mutants occur in the SF3 complex, which consists of the SF3a and SF3b subcomplexes (Brosi et al., 1993). Recent work has shown that the HEAT-repeat region of SF3B1, a SF3b component, undergoes a conformational transition upon U2 snRNP binding to the branchpoint, moving from an open conformation to a closed state, thereby stabilizing the U2-branchpoint duplex (Tholen et al., 2022; Zhang et al., 2021; Zhang et al., 2020a). We mapped an SF3B3 E159K mutant to an interface between SF3B3, SF3B1 and PHF5A (Figure 2D), distal to where PB or the branchpoint engages SF3b. The mutation lies in a region of SF3b that changes conformation upon binding to the branch helix (formed between U2 and the branchpoint sequence) (Figure 2E), likely impacting SF3B1 closing which would favor PB binding and, presumably, cell growth inhibition. Finally, we mapped PB-sensitive substitutions in residues in several factors not part of SF3b onto available structures (Haselbach et al., 2018; Nameki et al., 2022; Zhang et al., 2018) and found that they often occur at protein-protein interfaces (Figure S3D-F, Table S2), providing a resource for structure-functional studies.

### Viable SF3b mutations produce resistance to pladienolide B

SF3B1 and PHF5A mutations have been identified that render cells resistant to PB treatment (Cretu et al., 2018b; Teng et al., 2017a), but they only occur in a dominant (heterozygous) fashion, suggesting recessive lethality. Nonetheless, in our haploid screen, we identified eleven significantly enriched sgRNAs that target these factors, five against *PHF5A* and one against *SF3B1,* (Figure 1F). To identify the underlying alleles, we transduced eHAP FNLS cells with individual sgRNAs and collected cells at t0, t8 and t15 in the presence and absence of 2 nM PB treatment. Following genomic DNA isolation and amplicon deep sequencing, we determined mutation prevalence across time and treatments (Figure 3A). Note that for efficient guides, resistance-promoting mutations may be highly prevalent at t0 and therefore may not enrich in abundance under compound treatment (see e.g., sgPHF5A_7, Figure S3A).

**Figure 3.**
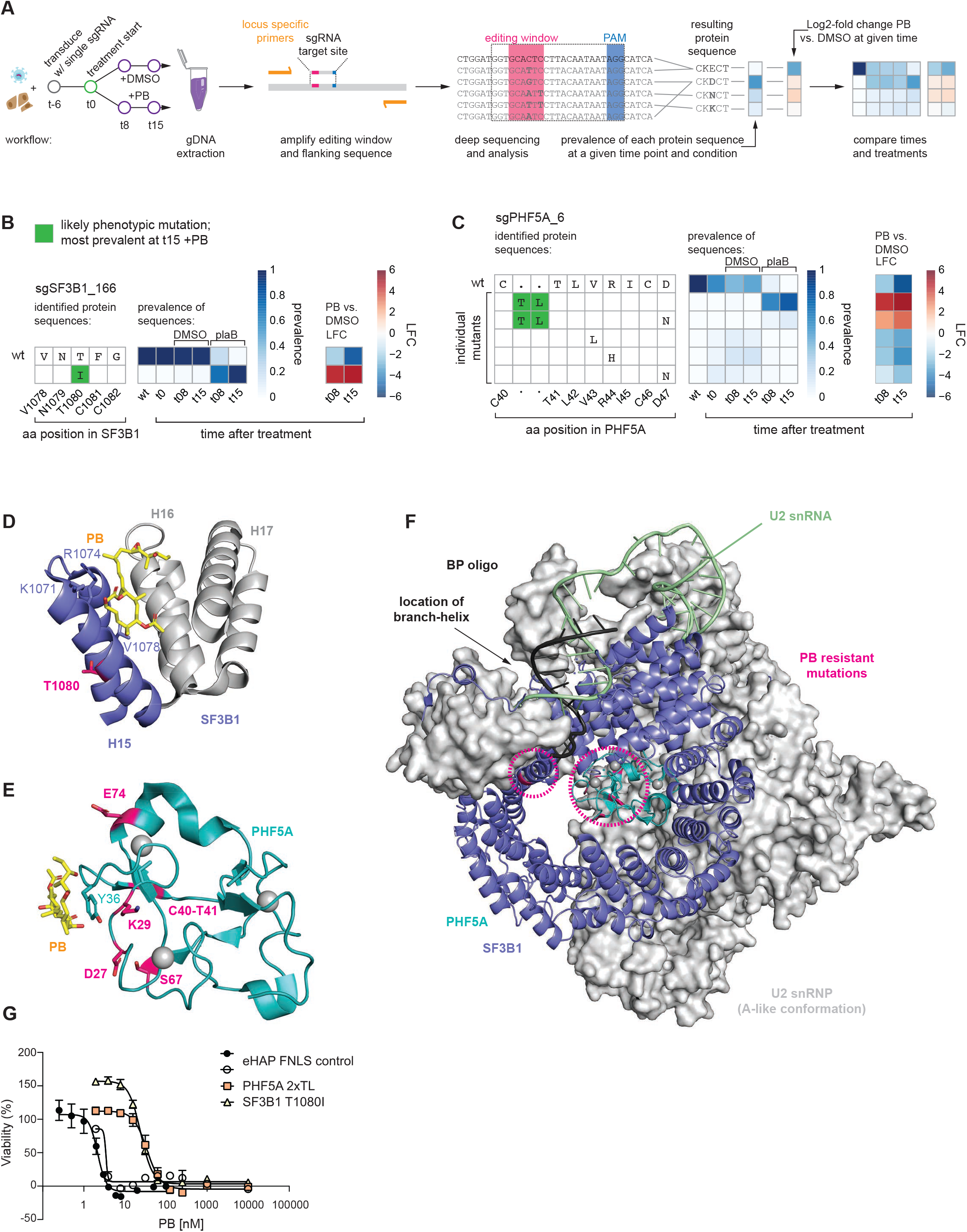
Novel resistance mutations in SF3B. (A) Schematic of workflow to identify phenotypic mutations. *Left:* Cells are transduced with single sgRNA in an arrayed format. After six days (t0) treatment is initiated and a cell sample is harvested at t0 as well as t8 and t15 for genomic DNA (gDNA) extraction. *Middle:* Locus-specific primers are used to amplify the editing window and its flanking sequence of the gDNA. The amplicons are then deep sequenced to identify mutations. *Right:* Mutational outcomes are translated and the resulting protein sequences are aggregated as multiple DNA sequences may result in the same protein sequence. Prevalence of each protein sequence is calculated for each time point and treatment condition. Where applicable, log2-fold changes are calculated between two samples. Finally, time points and treatments can be compared across samples for both prevalence and/or log_2_-fold change to the wild type (wt). Inferred phenotypic mutations (most prevalent at t15 +PB) are indicated by a green background. (B) Editing outcome for sgSF3B1_166: In the absence of PB across t0, t08 and t15 the wt is the only occurring protein (100% prevalence). Upon PB treatment, SF3B1 T1080I rapidly enriches (t15 = 100% prevalence). This striking behaviour is emphasised by the log_2_-fold change for this sgRNA. (C) Editing outcome for sgPHF5A_6: In the absence of PB some mutations occur within the editing window around R44. PB treatment enriches for a rare 6 nucleotide insertion occurring from the nicking action of the nCas9, which is part of FNLS. The resulting mutation is PHF5A TL insertion between C40 and T41 in PHF5A resulting in a tandem TLTL sequence. (D) Location of SF3B1 T1080I resistance mutation: SF3B1 HEAT repeats 15,16, and 17 (H15, H16, H17), which form part of the PB binding pocket (PB illustrated in yellow). H15 and H16 form a hinge within SF3B1 which undergoes a closing motion upon BP-A binding. T1080 (magenta) is located on the back of H15 facing away from PB and towards H14 (not shown). Known resistance mutations at K1071, R1074, V1078 are shown with side chains indicated. (PDB: 6EN4) (E) Location of PHF5A resistance mutations: All mutations are indicated (magenta) and occur on the face of PHF5A involved in PB binding. Y36C is also indicated – the only previously known PB resistance mutation in PHF5A. (PDB: 6EN4) (F) Illustration of SF3B1 and PHF5A resistance mutations in context of U2snRNP (A-like conformation). Mutations (magenta, circled) occur in vicinity to the branch helix and branchpoint adenosine. (PDB: 7Q4O) (G) Sixty-hour cell proliferation profiling (CellTiter-AQueous cellular viability and cytotoxicity assay) of control eHAP FNLS cell line expressing non-targeting sgRNA and monoclonal cell lines carrying either SF3B1 T1080I or PHF5A 2xTL mutation to PB. Error bars indicate s.d. n = 3 (average of two technical replicates for independent clonal cell lines).

sgSF3B1_166 was the only sgRNA conferring PB resistance through mutation of SF3B1. For this guide, no mutant alleles were detected by sequencing in the transduced cell population in the absence of PB treatment (possibly due to poor editing efficiency), but an allele encoding a T1080I change enriched rapidly upon PB addition to cells carrying this sgRNA (Figure 3B). In contrast, all five guides targeting *PHF5A* gave rise to substantial cell populations carrying different mutations at t0. Both sgPHF5A_7 and sgPHF5A_21 are predicted to result in cysteine to tyrosine mutations for side chains involved in the coordination of a Zn^2+^ ion. However, in both instances deep sequencing revealed that the predicted Cys > Tyr mutation was not detected; rather, we observed mutations impacting the preceding residue (Figure S3A,B). Both the resulting PHF5A K29N and E74D mutation arise through transversion (C > R mutation). sgPHF5A_47 is predicted to result in mutation and loss of the 3’ splice site of exon 3. This mutation did occur at high frequency (Figure S2C) and likely resulted in a growth disadvantage (see Figure S2A, reduced fitness of sgPHF5A_47 transduced cells in absence of PB). Instead, after transduction with this guide, the more frequent mutation at t0 encodes a D27N change which further accumulates in the population under compound selection. The remaining two resistance-promoting sgRNAs in PHF5A are predicted to be non-editing. sgRNA_PHF5A_6 targets a cytosine at position 7 in a G_6_C_7_ dinucleotide context, which is unfavourable to editing (Kluesner et al., 2021; Sánchez-Rivera et al., 2022). Indeed, this sgRNA resulted only in 12% of amplified molecules harboring the anticipated D47N mutation. Surprisingly, under PB selection, the more frequent mutation encodes a two amino acid insertion (TL) between C40 and T41 producing a tandem TL dipeptide in the protein sequence (hence we name the allele PHF5A-2xTL) (Figure 3C). This TL-encoding insertion occurred at position −3 relative to the PAM of sgRNA_PHF5A_6 where the nCas9 of the CRISPR-Cas9 base editor nicks the genomic DNA upon genomic binding. For the other non-editing sgRNA, sgRNA_PHF5A_26, editing should not occur within the editing window due to an absence of cytosines. Indeed, we observed edits 13 and 15 bp upstream of the targeted sequence, which alter S67 (Figure S3A, see Figure S3E for a summarized comparison of predicted mutations vs those identified by sequencing).

Using this information, we mapped the encoded amino acid changes on the available structures of SF3B1 and PHF5A. For SF3B1, the change lies within heat repeats 15 and 16, which form a hinge region that supports a conformational change necessary for BP-A binding (Figure 3D,F) (Cretu et al., 2018a; Tholen et al., 2022; Zhang et al., 2020b). This location differs from those of reported resistance mutations in SF3B1 at K1071, R1074 and V1078 – which are all residues that face PB in high resolution structures (Teng et al., 2017b; Yokoi et al., 2011). For PHF5A, all five mutations that we identified impact residues in a protein surface near the PB binding site, where PHF5A interacts with both SF3B1 and the U2-branchpoint helix (Figure 3E,F) (Tholen et al., 2022) This contrasts with the location of the reported (dominant resistant) Y36C mutation, which changes a residue that directly contacts PB (Teng et al., 2017b).

To test whether these mutations produce resistance to concentration-dependent acute killing by PB, we repeated guide transduction experiments followed by single cell cloning to generate six independent monoclonal cell lines, three harboring the mutation PHF5A-2xTL and three harboring SF3B1-T1080I (Figure S4A). The measured half maximal effective concentration (EC_50_) of PB in a cell viability assay at 60 h post treatment confirms that PHF5A-2xTL (34 nM) and SF3B1-T1080I (24 nM) confer concentration-dependent resistance to killing by PB relative to the parental cell line (EC_50_ = 2 nM) (Figure 3G).

### Mutations in the G-patch tumor suppressor protein SUGP1 confer PB resistance

Unexpectedly, our screen identified resistance mutations in three factors, PRPF6, SF1 and SUGP1, that are not part of SF3b, the target of PB. We focussed on the analysis of the SUGP1 mutations. *SUGP1* (SURP and G-patch domain containing 1) was targeted by two guides, both of which were confirmed in competition validation assays (Figure 1F, 2B, S2A). Transduction, compound treatment, and amplicon deep sequencing suggests that sgSUGP1_238 produces resistance via an E554K mutation and sgSUGP1_188 produces resistance via a G603N mutation (Figure 3B, C).

Both mutations match the editing predictions with one encoding a change lying in (G603N) and the other just upstream (E554K) of the G-patch motif (Figure 3A,D). SUGP1 is associates with the spliceosomal A complex, where it interacts with SF3B1 (Zhang et al., 2019). SUGP1 has not yet been visualized in any spliceosome structure, nor are there orthologs in *S. cerevisiae* or *S. pombe.* Recent work has identified SUGP1 as a putative tumor suppressor: its loss from the spliceosome was suggested to underlie splicing and oncogenic phenotypes of SF3B1 tumor mutations, and mutations in SUGP1 found in tumors mimic the splicing phenotype of SF3B1 mutant tumors (Alsafadi et al., 2021; Liu et al., 2020; Zhang et al., 2019).

For EC_50_ assays, we again repeated transductions and produced three independent monoclonal cell lines (Figure S4A) for each mutation. To our surprise, no substantial change in EC_50_ for PB was observed (Figure S4B). However, the EC_50_ measurement occurs over a much shorter time frame than treatment during the screen, which indicates that the mutations confer resistance to PB-mediated growth inhibition over time but not immediately. These data suggest that the SUGP1 mutants act by a mechanism distinct from those in SF3B1 and PHF5A, which map near the drug binding site.

### SUGP1 mutations modulate a subset of PB-induced exon skipping events

PB induces massive exon skipping as well as other splicing changes (Wu et al., 2018). To investigate the impact of SUGP1 mutations on PB-induced splicing changes, we performed RNA-seq analysis on our clonal cell lines. We also included the PHF5A-2xTL and SF3B1-T1080 clonal cell lines. Cells were treated with DMSO or 2 nM PB (= EC_50_) for 3 h (a time frame where cell viability is not yet affected, see Figure S4C) prior to RNA extraction and polyA-selection. We detected no changes in global transcript levels in the mutant cell lines (Figure S4C).

We used rMATS (Shen et al., 2014b) to detect differential alternative splicing events. In untreated cells, we observed exon skipping (skipped exon, SE) as the most frequent event triggered by the mutations followed by alternative 3’ splice site (A3’SS) use; relatively few introns were impacted [using a difference percent splicing inclusion (ΔPSI) cut-off of ≥ |10%| and FDR > 0.01] (Figure 4E). Among these were changes in 3’ splice site usage in introns of the *TMEM14C* and *ENOSF1* genes, which, strikingly, correspond exactly to changes observed previously in cells harboring SF3B1 cancer mutations or SUGP1 cancer mutations (Figure 4F, G, S4D-F) (Alsafadi et al., 2021; Liu et al., 2020).

**Figure 4.**
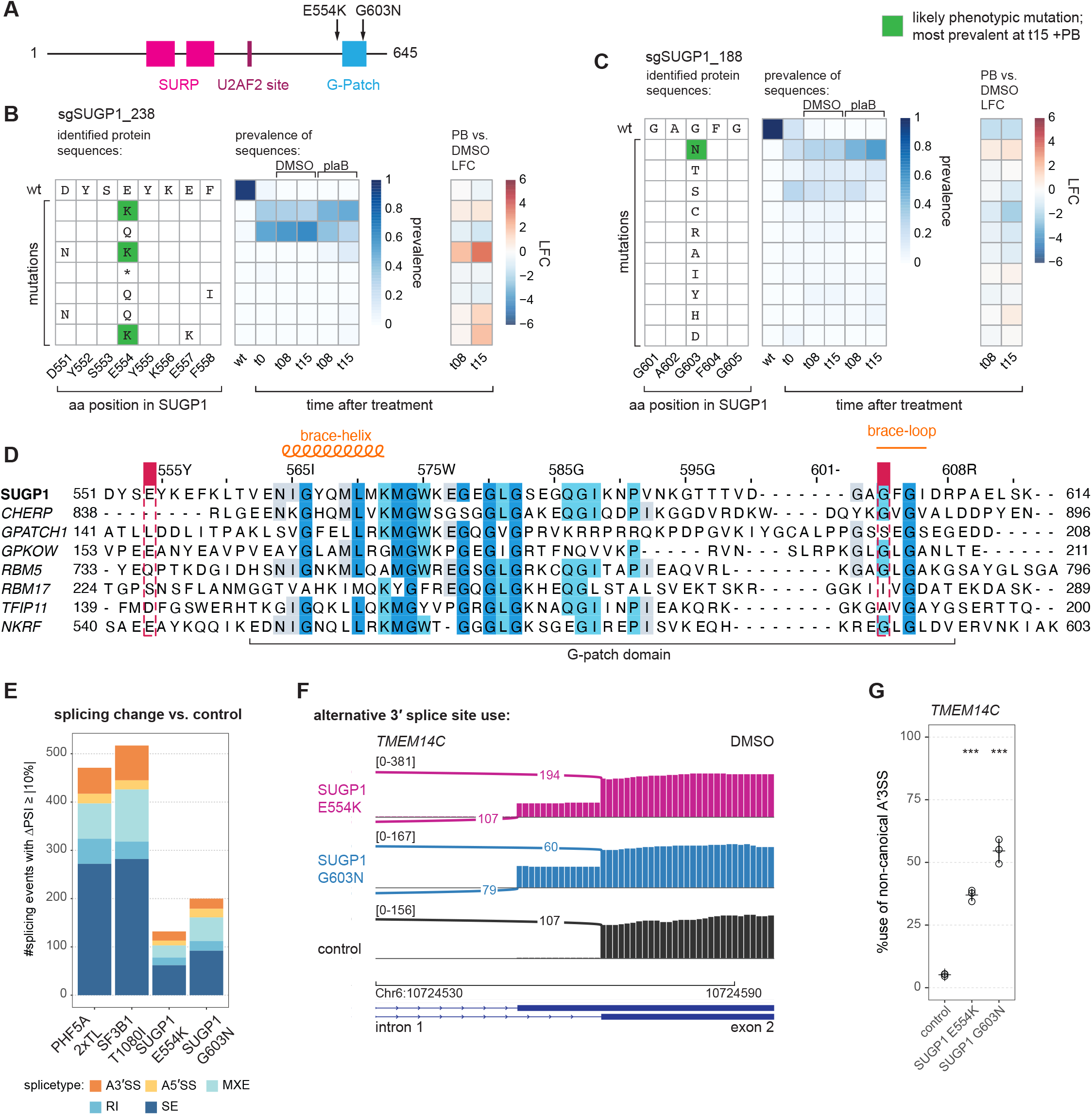
Novel resistance mutations in SUGP1. (A) Schematic of SUGP1 and its domains and motifs. Residues targeted and mutated by CRISPR-Cas9 base editor are indicated with arrows. (B) Editing outcome for sgSUGP1_238: At t0 and in the absence of PB treatment E554Q dominates. This mutation depletes under PB treatment and E554K becomes the dominant mutation. (C) Editing outcome for sgSUGP1_188: PB treatment enriches for G603N. Inferred phenotypic mutations are indicated with a green background and inferred bystander mutations with a yellow background. (D) Sequence alignment of all human G-patch motifs involved in splicing with the NKRF G-patch motif included as a reference for structural comparison. Shaded residues indicate amino acids with more than 30%identity. Brace-helix and brace-loop, as identified in the NKR G-patch are marked above the aligned sequences. Positions of mutants identified in screen are also indicated. Sequence alignment was performed with JalView (v.2.11.2.5). (E) Identified splicing changes for mutant vs. control cell lines. Skipped exons and A3’SS are the most frequently observed splicing changes. Numbers are shown for junctions identified with rMATS with FDR > 0.01 and |ΔPSI| ≥ 10 (ΔPSI: PSI of mutant sample - PSI of control sample, where PSI: percent spliced in). RNA-seq data of total, polyA-selected RNA from three independent clonal cell lines treated for 3 h with DMSO. (A3’SS: alternative 3’ splice site use; A5’SS: alternative 5’ splice site use; MXE: mutually exclusive exon; RI: retained intron; SE: skipped exon.) (F) Sashimi plot for alternative 3’ splice site usage in *TMEM14C* exon 2 (DMSO). The control cell line exclusively uses the distal 3’ SS while the SUGP1 mutants also make use of a proximal 3’SS. Representative traces for a single clonal cell line are shown (plot generated with IGV; alignment to Hg38). (G) RT-PCR and quantification for alternative 3’ splice site usage for *TMEM14C* exon 2 (DMSO). RNA extracted from eHAP FNLS cell lines - same monoclonal cell lines as for RNA-seq. *Statistical analysis for RT-PCR:* one-way ANOVA with Dunnett’s for multiple comparison (two-sided, with control as reference) was performed with R and package multcomp (v.1.4-20); * p < 0.05, ** p < 0.01, and *** p <0.001; all with n = 3.

Upon PB treatment of these cell lines, the number of differentially spliced junctions increased drastically. Control cells (eHAP FNLS transduced with a non-targeting sgRNA) displayed the largest number of PB-induced splicing changes, while the PHF5A-2xTL and SF3B1-T1080 showed almost no changes, as would be expected if they were to reduce the effects of compound binding to the spliceosome (Figure 5A). Both SUGP1 mutants displayed an intermediate phenotype with fewer affected events than wild-type in the presence of PB.

**Figure 5.**
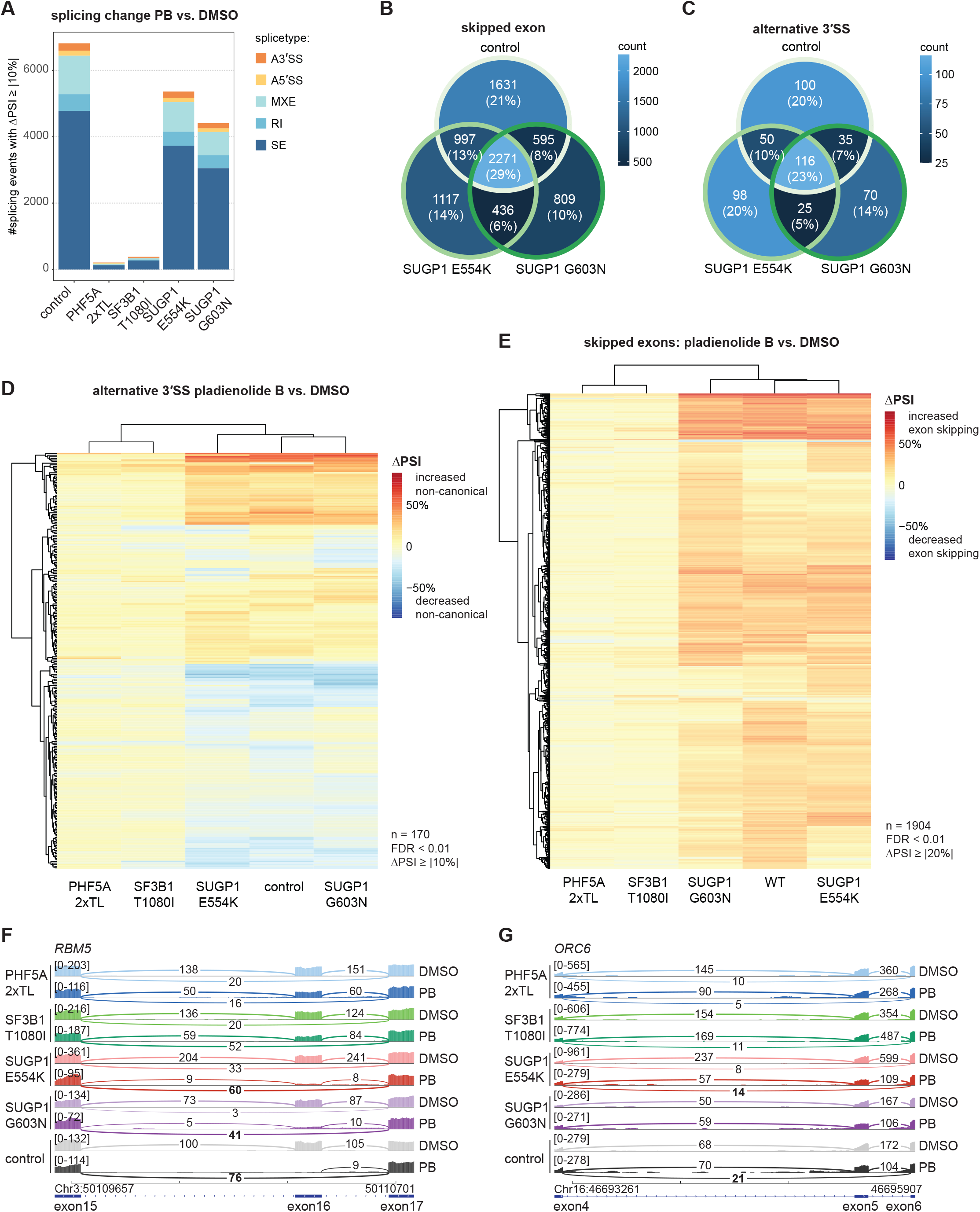
RNA-seq analysis of mutants. (A) Analysis of PB-induced splicing regulation in mutant vs. control cell lines. PB-resistant cell lines carrying mutations PHF5A 2xTL or SF3B1 T1080I at the PB binding pocket show very little change in splicing regulation compared to control cell lines or SUGP1 mutant cell lines. Numbers are shown for junctions identified with rMATS with FDR > 0.01 and |ΔPSI| ≥ 10. RNA-seq data of total, polyA-selected RNA from three independent clonal cell lines treated for 3 h with 2 nM PB or DMSO. (B) Overlap in cassette exons affected by PB treatment for all junctions observed in all three sample groups (control vs. SUGP1 E554K vs. SUGP1 G603N). 29% of differentially spliced cassette exons are affected in all three genetic backgrounds. SUGP1 mutants do not share more overlap than they individually share with the control sample. Splicing junctions had to be detected by rMATS with FDR < 0.01 to be included in analysis. (C) Overlap in alternative 3’ splice site use affected by PB treatment for all junctions observed in all three sample groups (control vs. SUGP1 E554K vs. SUGP1 G603N). 23% of differentially spliced cassette exons are affected in all three genetic backgrounds. SUGP1 mutants do not share more overlap than they individually share with the control sample. Splicing junctions had to be detected by rMATS with FDR < 0.01 to be included in analysis. (D) Hierarchical clustering of differential splicing of alternative 3’ splice sites for PB treatment, based on PSI (percent spliced in) changes. The heatmap represents ΔPSI values of A3’SS use upon treatment with PB at 2 nM for 3 h vs. DMSO as detected in total, polyA-selected RNA using rMATS. In contrast to mutants at the PB-binding interface, SUGP1 mutants E554K and G603N are susceptible to PB induced splicing changes. Predominantly, PB treatment results in increased and decreased use of the canonical 3’SS. Mutations in SUGP1 modulate observed phenotype. Only splicing junctions with FDR < 0.01 and |ΔPSI| ≥ 10 were considered. (E) Hierarchical clustering of differential splicing of cassette exons for PB treatment. The heatmap represents ΔPSI values of cassette exons upon treatment with PB at 2 nM for 3 h vs. DMSO as detected in total, polyA-selected RNA from eHAP FNLS cells using rMATS. In contrast to mutants at the PB-binding interface, SUGP1 mutants E554K and G603N are susceptible to PB induced splicing changes. Predominantly, PB treatment results in increased exon skipping and mutations in SUGP1 modulate observed phenotype. Only splicing junctions with FDR < 0.01 and |ΔPSI| ≥ 20 were considered. (F) Sashimi plot for alternative splicing of exon 16 in *RBM5* for 2 nM PB vs. DMSO treatment. The control cell line and the SUGP1 mutants switch to predominant exon skipping upon PB treatment. Representative traces for a single cell line each (plot generated with IGV; alignment to Hg38). (G) Sashimi plot for alternative splicing of exon 4 in *ORC6* for 2 nM PB vs. DMSO treatment. SUGP1 mutants show less exon skipping than control cell lines upon PB treatment. Representative traces for a single cell line each are shown (plot generated with IGV; alignment to Hg38).

Focusing on A3’SS and SE events induced by PB, we observed a general concordance between control cells and SUGP1 mutants (Figure 5B, C). Hierarchical clustering of the splicing junctions quantified by rMATS across all samples demonstrated that PB treatment almost exclusively triggered increases in exon skipping (Figure 5E), but both increases and decreases use of alternative 3’ splice sites (Figure 5D). Subsets of splicing events induced by PB were affected either equally strongly in control and SUGP1 mutant lines (example: *RBM5* exon 16 in Figure 5F, S5B) or displayed milder changes in the SUGP1 mutants (example: *ORC6* exon 5 in Figure 5G, S5B, C). No statistically significant differences in intronic or exonic features at the introns equally vs. differentially affected by SUGP1 genotype were identified using Matt (Gohr and Irimia, 2019); thus additional work will be necessary to identify the determinants of SUGP1-sensitive, PB-induced alternative splicing events. These findings demonstrate that SUGP1 mutants can modulate splicing changes induced by PB, as expected from the ability of SUGP1 to produce relative resistance to the compound.

### DHX15/hPrp43 is a mutationally sensitive ligand of the G-patch motif of SUGP1

G-patch motifs (named for their glycine-richness) are direct activators of DEAH-box helicases (Studer et al., 2020a; Warkocki et al., 2015); this is the only known activity of this domain. SUGP1 has therefore been suggested to recruit a helicase through its G-patch motif to SF3B1 and the A complex (Alsafadi et al., 2021; Liu et al., 2020; Zhang et al., 2019). As we identified mutations within and just upstream of the SUGP1 G-patch, it seemed likely that one or both mutations impact the association and/or activation of a cognate DEAH-box ATPase. Indeed, the G603N mutation lies within the “brace loop” analogous to the only G-patch protein for which its helicase-bound structure is available (Figure 4D), a region known to be important for the NKRF G-patch to bind and activate its cognate DEAH box helicase DHX15 (Studer et al., 2020b).

To identify SUGP1 helicase partner(s), we employed proximity labelling (BioID) exploiting our SUGP1 G603N mutant as a control. Recent advances allow for short labelling times using Turbo fusion proteins (Branon et al., 2018). After optimizing conditions, we transfected HEK293T cells with expression plasmids encoding SUGP1 with a C-terminal miniTuboID fusion. After a short 2 min labelling pulse with biotin, extracts were prepared, and labelled proteins purified under harsh conditions using streptavidin. Replicate samples were subjected to tandem mass tag mass spectrometry (TMT-MS) (Figure S6A,B) to identify differentially labelled proteins. Remarkably, only a single protein displayed a statistically significant reduction in signal in the SUGP1-G603N-miniTurboID-tagged samples versus those obtained with the wild-type fusion: the DEAH-box helicase DHX15/hPrp43, the protein that disassembles the spliceosome (Figure 6A). We also identified DHX15/hPrp43 in a parallel experiment using a fusion of a SUGP1-L570E-miniTurboID fusion which we constructed based on the ability of this mutation to disrupt G-patch/DEAH protein interactions for a different G-patch protein, NKRF (Studer et al., 2020b) (Figure S6C).

**Figure 6.**
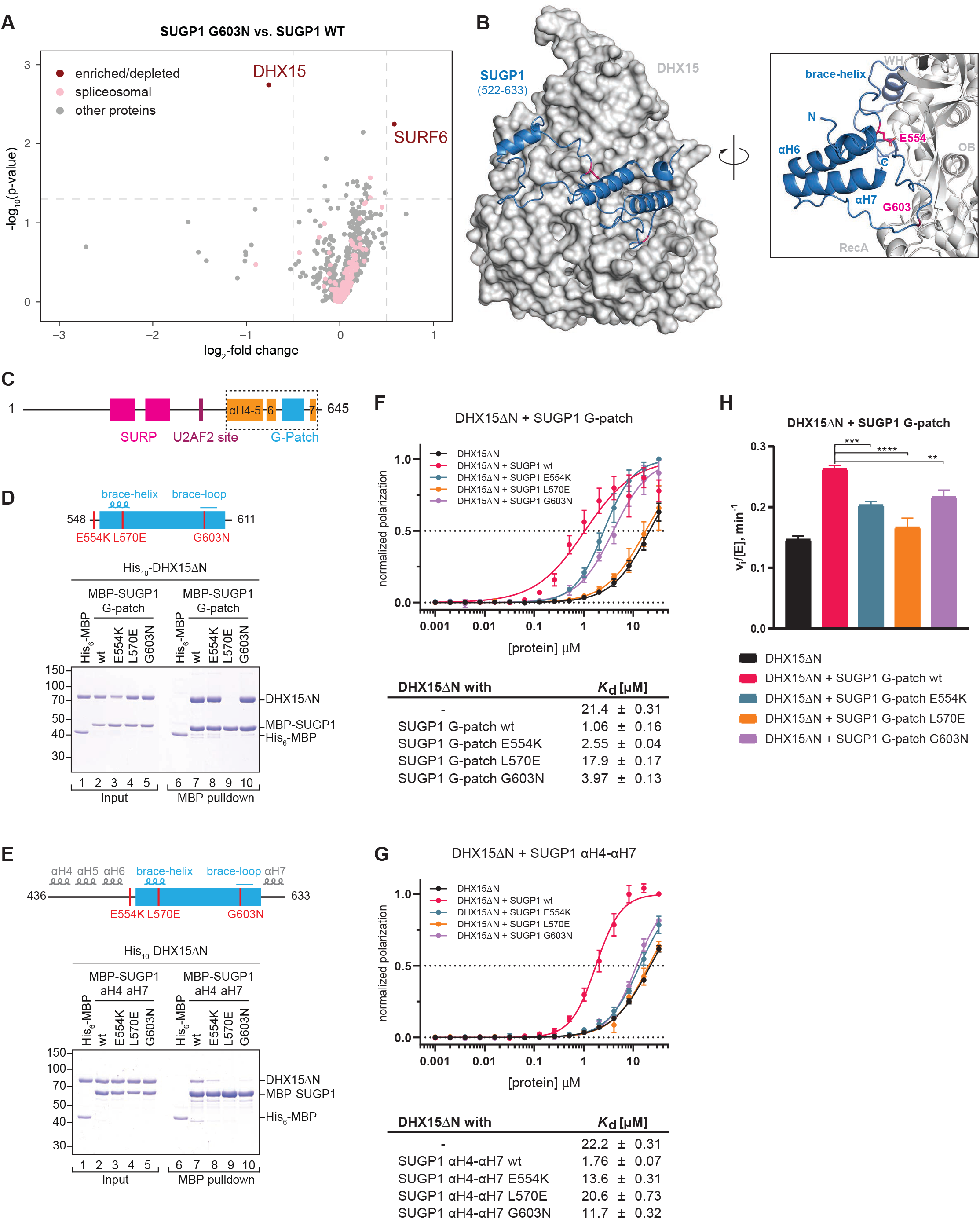
SUGP1 interacts with DHX15. (A) miniTurbo proximity labeling using FLAG-SUGP1-miniTurboID G603N vs. wt overexpressed in HEK293T cells. Biotinylated proteins were identified with TMT-MS after enrichment with streptavidin beads. −log_10_(p-value) is plotted against the log_2_-fold change (LFC) in a volcano plot (dashed lines: cutoffs at p < 0.05 and LFC > 0.5). The only identified depleted factor is DHX15. SURF6 is a protein of the ribosomal biogenesis pathway. (B) AlphaFold2 prediction of SUGP1 (522-633), encompassing the G-patch motif flanked by αH6 and αH7, in complex with DHX15. To the right a close up is shown. The residues identified in our screen (E554 & G603, magenta) are at the interface with DHX15, while αH6 & αH7 are not predicted to interact with the helicase. (n = 4) (C-E) Domain organization and schematic representation of the MBP-*hs*SUGP1 variants with the introduced mutations colored in red. The different SUGP1 construct boundaries for G-patch (amino acid residues 548-611), αH6-αH7 (522-633) and αH4-αH7 (436-633), respectively, are indicated at the sides and features of the predicted secondary structure are depicted above. (D-E) Coomassie-stained gels of protein binding assays using purified MBP-*hs*SUGP1 constructs and His_10_-*hs*DHX15ΔN. MBP-SUGP1 G-patch (D), and αH4-αH7 (E) with either wildtype (wt) protein sequence or carrying the indicated mutation were used as baits and His_6_-MBP served as a control. Input (1.5% of total) and eluates (24% of total) were loaded. (F-G) Fluorescence polarization of FAM-labeled U_12_ RNA with His_10_-*hs*DHX15ΔN in the absence or presence of MBP-SUGP1 G-patch (F) and αH4-αH7 (G) wt or mutants. The dashed line indicates 50% normalized polarization. Error bars represent standard deviations from the average values of triplicate measurements. RNA dissociation constants (Kd) with standard error of means (SEM) were derived from fitting the respective data by linear regression. (H) Initial ATPase activity rates of His_10_-*hs*DHX15ΔN in the absence or presence of MBP-*hs*SUGP1 G-patch wt or mutants at 250 μM ATP. Error bars indicate standard deviations of three independent measurements, asterisks denote significance (one-way ANOVA with Tukey’s) with ** p < 0.01, *** p < 0.001, and **** p < 0.0001.

To test more directly the impact of the G-patch mutations on the interaction and/or activation of with DHX15, we employed biochemical methods. Modelling of a SUGP1-DHX15 using AlphaFold2 (Jumper et al., 2021) predicted that the G-patch domain of SUGP1 interacted with DHX15 as anticipated but that the G-patch was flanked by unanticipated α-helical elements that are also predicted to interact with DHX15. In this model, the E554K substitution may impact a contact with the flanking α-helices (Figure 6B). We overexpressed SUGP1 mutants in HEK293T cells to assess their impact on splicing. We observed that deletion of αH4-5 or αH6 as well as the E554K, G603N, and L570E substitutions displayed similar effects on splicing (Figure S5D-G. Thus, we constructed a series of maltose binding protein (MBP) fusion proteins harboring the SUGP1 G-patch and varying lengths of flanking sequences (Figures 6D,E and S6C).

We next mixed a purified version of DHX15 lacking its N-terminal domain (DHX15ΔN) with each MBP-SUGP1 fusion proteins, purified them with amylose beads, and analyzed the material using SDS-PAGE. We observed that DHX15ΔN selectively copurified with each of the MBP-SUGP1 fusion proteins (Figure 6D-E and S6D). However, we found that the efficiency of copurification decreased with increasing length of SUGP1 constructs, indicating that protein stretches surrounding the G-patch modulate its binding affinity for DHX15. For each of these constructs, we generated mutations in the G-patch, corresponding to the two obtained in our screen, E554K and G603N, as well as one in the central “brace-helix”, L570E, that is known to disrupt ligand binding in analogous G-patch proteins. For the construct harboring the most upstream SUGP1 sequences (436-633) all mutations reduced binding to DHX15ΔN, consistent with loss of DHX15 labelling we observed in the BioID experiment. Therefore, it is likely that disruption of DHX15 interaction forms the molecular basis for the observed PB resistance of the SUGP1 mutants obtained in our screen. Consistent with the stronger affinity of DHX15ΔN for the intermediate [MBP-SUGP1(522-633)] and shortest [MBP-SUGP1 (548-611)] SUGP1 constructs, their binding was not sensitive to the mutations obtained in our screen but was only disrupted by mutation of the core interface residue L570E.

As G-patch proteins can enhance RNA binding to DEAH-box helicases (Studer et al., 2020b), we used fluorescence anisotropy to ask whether the SUGP1 fusions increased RNA affinity of DHX15ΔN. Using a fluorescently labelled poly-U RNA, we found that all SUGP1 constructs indeed increased RNA binding to DHX15ΔN ~10-20-fold (Figure 6F-G and S6F). In agreement with their effect on binding DHX15, all mutations in the longest SUGP1 construct (436-633) blocked this stimulation (Figure 6G), while varying degrees of mutational sensitivity were observed for the shorter SUGP1 truncations (Figure 6G and S6G), again indicating an important role for sequences flanking the G-patch domain.

Finally, we investigated the ability of SUGP1 to stimulate the ATPase activity of DHX15ΔN that is required for and coupled to its RNP remodelling activity. Indeed, SUGP1 (548-611) produced a 1.8-fold increase in initial ATPase rates of DHX15ΔN at saturating ATP concentrations, which was significantly diminished by all mutations consistent with their negative effects on DHX15 binding.

## DISCUSSION

The human spliceosome is essential for the splicing of over 200,000 introns in the human genome. Because it is mutated in numerous diseases and the target of myriad splicing regulators, it is a key compound development target. Numerous questions exist regarding the coupling of transcription and chromatin to splicing, the underpinnings of splicing fidelity, and the functional roles of many if not most human spliceosomal proteins. However, the genetic analysis of the human spliceosome has not been pursued, even though it harbors ~60 proteins not found in *S. cerevisiae* and is likely to operate in ways that cannot be anticipated from prior studies of yeast. Particularly useful would be so-called “informative alleles” that dissect essential protein function. While CRISPR-Cas9 knockout screens are not designed to generate such information, CRISPR base editing and related methods in principle provide approaches generating programmed point mutations. We adapted pooled CRISPR-Cas9 base editing to the human spliceosome to mutagenize 153 protein subunits in a haploid cell context. We interrogated the mutants with PB, a prototype for a class of anti-cancer compounds that targets the SF3b complex. Our studies provide insights into structure-function relationships by identifying viable alleles of numerous spliceosomal proteins that program hypersensitivity or resistance to SF3b inhibition. We demonstrate the utility of such alleles through studies of the SUGP1 tumor suppressor. Below we discuss the evidence for these conclusions and propose a new human-specific discard/fidelity step mediated by the activation of the spliceosomal disassemblase DHX15/hPrp43 by SUGP1 during early stages of spliceosome assembly and its implications for human disease.

### PB hypersensitive mutations identify functional sites in the human spliceosome that vary in the human population

We obtained PB-hypersensitive mutants in a small subset of the 153 proteins mutagenized. Gratifyingly, most lie in factors that act at or near the step inhibited by PB, including in SF1 and components of U2 snRNP. This specificity highlights the utility of single-residue chemical-genetic interactions to identify functional sites related to a particular phase of an essential process. As described in the results above, many mutations we identified could be placed on existing structures, enabling the generation of structure-function relationships. However, most of the residues altered in our mutants are not visualized in existing structures. In both cases, detailed studies *in vitro* and/or *in vivo* will be required to understand the impact of these functional sites on splicing of the numerous endogenous introns that interrupt human genes. The mutations identified here provide a resource for such investigations.

Mutations in the second-step factor CDC40/hPrp17 and CACTIN produce PB hypersensitivity to PB, even though PB impacts SF3b, which is dissociated from the spliceosome (freeing the U2-branchpoint helix) by DHX16/Prp2 prior to the chemical steps of splicing so that the active site of the spliceosome can form. We speculate that triggering the use of different branchpoints via PB results in a dependency on weaker 3’ splice sites, whose docking into the catalytic core of the spliceosome requires stabilization by step 2 factors, a model consistent with the cryoEM structure of the human post-catalytic P complex (Fica et al., 2019).

Because the residues impacted by the PB-sensitive mutations are (by definition) functional, one anticipates that they would not vary in the human population, given the essential function of the spliceosome. Nonetheless, we asked this question by searching the ClinGen database. Strikingly, residues altered in seven of the PB-sensitive mutants have been identified in the human population (labelled as “non-disease-associated”) with four displaying the exact same amino acid changes in the human population as in our PB-sensitive cells (Table S2), suggesting that these variants very likely have a functional impact.

### Identification of SUGP1 G-patch mutations as PB-resistant

Our studies identified two sgRNAs that target SUGP1 that produce PB-resistance. The stronger of the two alleles produced by these guides, G603N, lies in a conserved domain of G-patch proteins called the “brace loop” which is important for G-patch proteins to activate their cognate DEAH-box helicase. The other change, E554K, lies just upstream in a region predicted by AlphaFold2 to be helical. By performing proximity labelling, comparing wild-type versus mutant proteins, we identified a single DEAH-box protein, DHX15/hPrp43 as being both labelled by SUGP1-miniTurboID fusions and sensitive to a G-patch mutation obtained in our screen. DHX15 has often been found in proteomic studies of early spliceosomes, but its function at this stage remained relatively opaque. However, a recent study from Jurica and colleagues has shown that depletion of DHX15 from HeLa cell extracts results in an increase rather than a decrease in A complex formation (Maul-Newby et al., 2022). This result is consistent with the observation that yeast Prp43 is the helicase that disassembles spliceosomes (Martin et al., 2002; Tanaka et al., 2007). Taken together with our results and the known association of SUGP1 with early spliceosomal complexes, we propose that SUGP1 recruits DHX15 to disassemble early spliceosomes, constituting an early discard step analogous to late discard steps described by Staley and colleagues in yeast (Figure 7). In this model, PB resistance results from mutation in the SUGP1 G-patch domain because this increases A complex formation or residence time by inhibiting disassembly, thereby counteracting the inhibitory activity of PB in reducing stable A complex formation.

**Figure 7.**
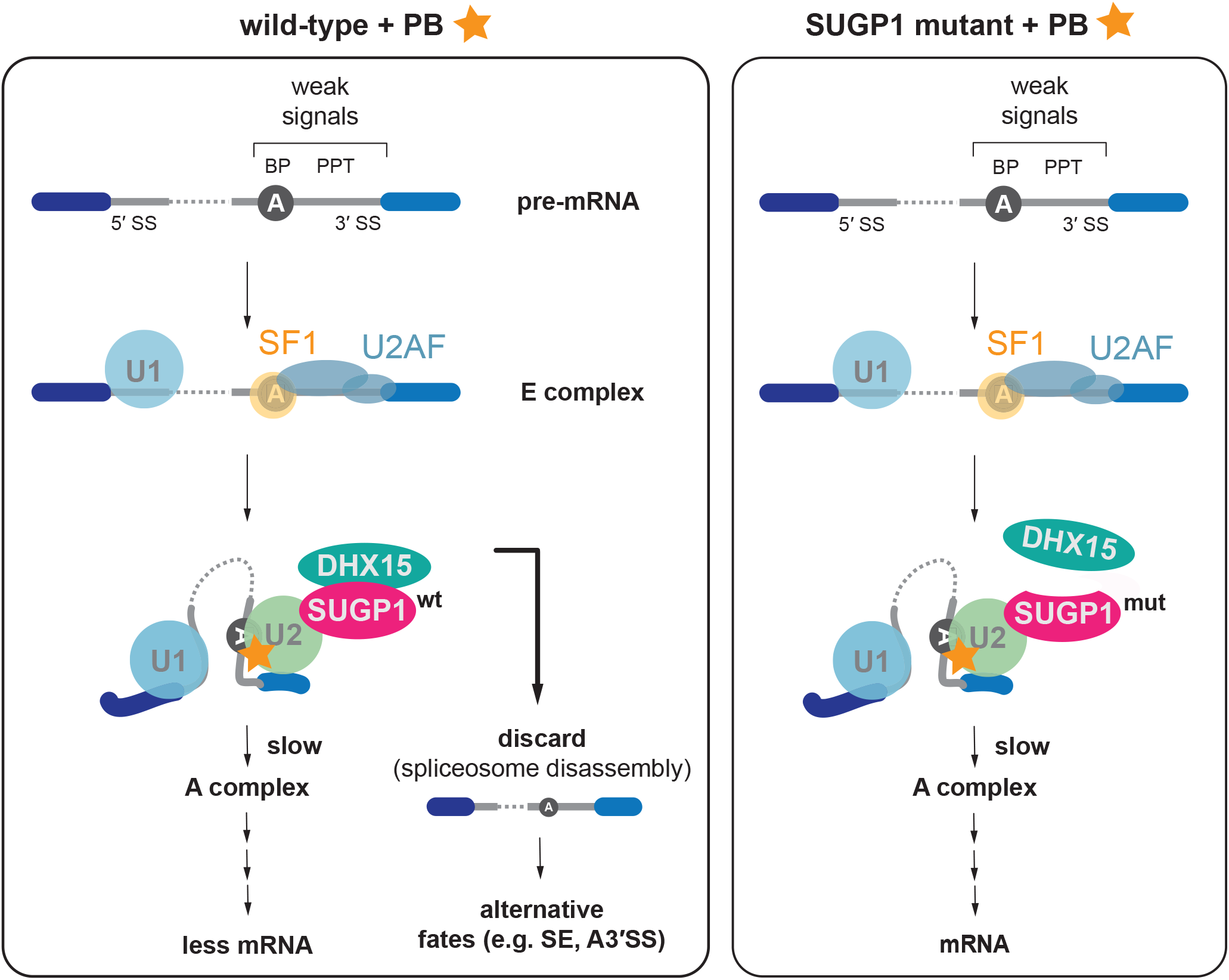
Model for SUGP1-DHX15 and proofreading at early spliceosome assembly. *Left panel:* On weak splice sites (either with a weak branchpoint sequence, PPT or 3’ SS or a combination thereof) the transition from E complex to A complex is inhibited as PB binds to the U2 snRNP and prevents the full binding of the branch helix and recognition of the BP-A. These stalled spliceosomes are recognized by SUGP1-DHX15 and are discarded. Less mRNA is being produced in this scenario and more alternative mRNAs with skipped exons or alternative 3’ splice site choice result. *Right panel:* A mutation in SUGP1 can weaken SUGP1 interaction with DHX15, removing the proofreading & discard pathway, giving the cell “more time” to assembly A complex and to proceed with splicing.

### Relationship to oncogenic mutations in SUGP1 and SF3B1

Manley and colleagues have proposed that cancer SF3B1 mutations act by limiting association of SUGP1 with the spliceosome. This model, based on biochemistry, is supported by genetic data that identified SUGP1 mutations in tumors that mimic the splicing phenotypes of SF3B1 mutant tumors (Alsafadi et al., 2021). Indeed, many of the identified cancer mutations map to the regions flanking the G-patch motif of SUGP1 (Figure S5F), which we show to influence SUGP1-DHX15 interaction and splicing. It was proposed that the then-unknown helicase that is recruited by SUGP1 might dissociate SF1 from the branchpoint, causing U2 snRNP to relocate to alternative branchpoints (Zhang et al., 2019). However, our discard model proposes a different mechanism underpinning the effects of SUGP1 cancer mutations, namely a defect in an early rejection step mediated by spliceosome disassembly (Figure 7). Such a model would explain the activation of cryptic branchpoints as a defect in proofreading enabling the production of oncogenic mRNAs via the activation of cryptic branchpoint/3’ splice site combinations as has been observed in SF3B1 and SUGP1 mutants (Liu et al., 2020; Zhang et al., 2019). This model is also consistent with *in vitro* studies of DHX15 depletion described above and the known activity of DHX15 in spliceosome disassembly.

### Mutagenesis of an essential machine in human cells in a haploid context

While this work was underway, two laboratories recently independently reported the deployment of base editor libraries in a variety of screens that involved phenotypic characterization of single nucleotide variants and/or to probing of small molecule-protein interactions (Cuella-Martin et al., 2021; Hanna et al., 2021). These studies largely interrogated individual proteins or part of a gene network and were mostly performed in diploid (or diploid-like) cells. To enable large scale studies in a haploid context, we generated editing-competent eHAP cells expressing the FLNS editor under conditions that maintained their haploid state. A notable finding from this work is that editors produce unexpected mutations at low frequencies which can produce phenotypes upon positive selection. For PHF5A, about half of the PB-resistant mutants appeared to select for such mutations. While undesirable for the use of base editing to generate specific programmed alleles, such unanticipated activities are useful for mutagenesis studies. Given our experience with the spliceosome, the future is bright for using base editing to interrogate essential cellular machines in human cells to produce new insights into human cell biology and disease.

### Limitations of this study

The editor we used in this study targets only a subset of residues in spliceosomal proteins and is limited by base editor specificity and the occurrence of PAM sites. Thus, many additional potential informative mutations may be isolatable with the advent of complementary technologies that enable for efficient base editing at additional sites without causing unwanted cellular toxicity. Our analysis of PB-hypersensitive sites requires further studies to understand their mechanistic impact. Biochemical tests of the proofreading model will require its reconstitution *in vitro* and, ultimately, structural analysis.

## Supporting information

Suppemental Table S1

Suppemental Table S2

Suppemental Table S3

## Acknowledgements

We dedicate this paper to the memory of Dr. Christine Guthrie. We thank all members of the Madhani lab for scientific discussions and support. We thank Stefan Oberlin (UCSF) for advice on bioinformatics analysis and CRISPR screens, Lukas E. Dow (Cornell) and David R. Liu (Harvard) for advice on base editing, and Daniël van Leeuwen and Chris Richards for advice regarding HAP1 cells. The authors thank Qing Feng and Chris Burge for communicating unpublished results and for technical advice. Sequencing data was generated at the UCSF CAT and through the kind use of their NextSeq System by Venice Servellita and Charles Chiu. I.B. was supported by the Swiss National Science Foundation through an SNF Postdoctoral Fellowship (191929 and 203008). Research in the Madhani lab is supported by R01 GM71801. Mass Spectrometry was provided by the Mass Spectrometry Resource at UCSF (A.L. Burlingame, Director) supported by the Dr. Miriam and Sheldon G. Adelson Medical Research Foundation (AMRF) and the UCSF Program for Breakthrough Biomedical Research (PBBR). Work in the Jonas lab was supported by the Swiss National Science Foundation (SNSF) through the National Center for Excellence in Research “RNA & Disease” (grant number 205601) and through SNSF project grants (grant numbers 179498 and 207458).

## Author contributions

Conceptualization and Methodology, I.B. and H.D.M.; Validation, Formal Analysis, and Visualization, I.B.; Investigation, I.B., B.R., S.L., J.O-P., E.S., M.K.S, T. L., V.S.; Writing – Original Draft, I.B., S.J. and H.D.M.; Writing – Review & Editing, all authors; Supervision, I.B., H.D.M., and S.J.; Funding Acquisition, I.B., H.D.M and S.J.

## MATERIALS AND METHODS

### Lead contact

Further information and request for resources and reagents should be directed to and will be fulfilled by the Lead Contact, Hiten D. Madhani.

### Materials availability

Plasmids generated in this study are available from the Lead Contact. Cell lines generated in this study are not available as eHAP cells and any product derived thereof are protected under an MTA upon purchase of the eHAP parental cell lines from Horizon (original vendor).

### Data and code availability

The read counts for the CRISPR-Cas9 base editing screen is provided as Supplementary Tables and the FASTQ files for screens, validation experiments are deposited on the Sequence Read Archive. The accession number for the data reported is: XXXXX. RNA-seq data is deposited on the Sequence Read Archive with the accession number: XXXX. The mass spectrometry proteomics data have been deposited to the ProteomeXchange Consortium via the PRIDE (Perez-Riverol et al., 2022) partner repository with the dataset identifier PXD038067.

#### METHOD DETAILS

##### Vectors

Assembly of vectors (cloning and mutagenesis) was, if not otherwise indicated, performed using NEBuilder^®^ HiFi DNA Assembly (NEB #E2621).

###### pLibrary (MP783)

mU6 promoter expresses customizable guide RNA with a 20N barcode sequence at the 3’ end of the tracrRNA to facilitate identification of individual sgRNAs and sample splitting into replicates (Boettcher et al., 2019). A core EF1α promoter expresses puromycin resistance and a T2A site provides BFP for easy titer determination of the lentiviral library.

###### pFNLS

A core EF1α promoter expresses 3xFLAG-tagged codon optimized FNLS base editor and provides with a P2A site EGFP for identification of FNLS carrying cell lines. A PGK promoter provides blasticidin resistance. This vector was modified from Addgene vector #110869.

###### pHA-SUGP1

Point mutations and deletions were introduced with into p3xFLAG-CMV-14_3xHA (Zhang et al., 2019).

###### Vectors for BioID

SUGP1 constructs were cloned from pHA-SUGP1 using primers (IB0156 5’-atgacgtcccagactacgcagctagcAGTCTCAAGATGGACAACC-3’; IB0157 5’-tgtttagcgttcagcagcgggatagatccgcctgaGTAGTAAGGCCGTCTGG-3’) into pCDNA3_3xHA-miniTurbo-NLS (Addgene #107172) digested with NheI-HF (NEB #R3131).

###### Expression plasmids

SUGP1 constructs were generated by PCR using a plasmid from the human open reading frame library (hORFeome Version 5.1, ID: 53373) as template and gene-specific primers. For protein expression in E. coli, the constructs were cloned into the NdeI-XbaI sites of the plasmid pnEA-NpM, which is derived from the pET-MCN vector series that harbors an N-terminal MBP-tag and a subsequent 3C protease site (Haffke et al., 2015). Mutations in the SUGP1 constructs were introduced by ‘round-the-horn (RTH) mutagenesis (Hemsley et al., 1989) using the respective primers (Table S3). The cloning and insect cell expression of hsDHX15ΔN has been described previously (Studer et al., 2020b).

##### Cell Culture

All cell lines were maintained at 37 °C with 5% CO_2_ and were regularly tested negative for mycoplasma infection. Human embryonic kidney (HEK) 293T cells were grown in DMEM (with 4.5 g/L glucose, L-glutamine and sodium pyruvate; Corning #10-013-CV) supplemented with 10% FBS. Cells were passaged every 2-3 days.

Human eHAP cell lines and derivatives thereof were cultured in IMDM (with L-glutamine, with HEPES; Cytiva #SH30228), supplemented with 10% FBS and 1:100 penicillin/streptomycin. eHAP cell lines were at all times maintained at sub-confluent conditions and ploidy was regularly assessed with flow cytometry. When mentioned, doses of puromycin were 4 μg/ml and blasticidin 10 μg/ml.

Flow cytometry data was analysed with FlowJo (v10.8.1).

##### Cell viability assay

96-well plates were seeded with 11000 cells per well earlier in the day and treatment was started after allowing cells enough time to attach to the plate surface. A serial dilution of PB was then used (starting at 100 nM and followed by 10 additional 2-fold dilution steps down to 0.25 nM or starting at 10 μM, 1 μM, 250 nM and followed by 7 additional 2-fold dilution steps down to 0.98 nM). DMSO percentage was maintained throughout and a DMSO-only control was included. 60 h post PB addition, CellTiter 96^®^ AQ_ueous_ One Solution Cell Proliferation Assay reagent (Promega #G3582) was added and incubated for 4 h and read out according to the manufacturer’s instruction. Samples were measured in two technical replicates, whose values were used as an average for three biological replicates. EC_50_ curves were fit with GraphPad PRISM.

##### Spliceosome library design and production

We compiled a list of all spliceosome components reproducibly detected through mass spectrometry(MS), interaction studies, and/or purified and visualized in the spliceosome in structural biology studies (Sales-Lee et al., 2021). This list encompasses 153 proteins (Table S1). Guide sequences for targeting the spliceosome were designed using CHOPCHOP(Labun et al., 2019) using [-Target $GENE -J -BED -GenBank -G hg38 -filterGCmin 0 -filterGCmax 100 -consensusUnion -t CODING -n N -a 20 -T 1 -g 20 -M NGG]. We included all sgRNAs targeting coding sequence across all exons in all isoforms, including 20 nucleotides into the introns and UTR. Oligonucleotide pools were synthesised by CustomArray. Cloning sites were appended with 5’-AGTATCCCTTGGAGAACCACCTTGTTGG-3’ and 5’-GTTTAAGAGCTATGCTGGAAACAGCATA-3’. The final oligonucleotide sequence was thus: 5’-AGTATCCCTTGGAGAACCACCTTGTTGG [sgRNA, 20 nt] GTTTAAGAGCTATGCTGGAAACAGCATA-3’.

Primers (forward: cttggAGAACCACCTTGTTG, reverse: GTTTCCAGCATAGCTCTTAAAC) were used to amplify the library pool (15x cycles). The resulting amplicons were PCR purified (QIAGEN #28104) and cloned into the library vector [digested with AarI (ThermoFisher #ER1582)] via Gibson assembly (NEBuilder^®^ HiFi DNA Assembly). The ligation product was buffer exchanged (BioRad #732-622) ethanol precipitated and electroporated into MegaX DH10B T1^R^ Electrocomp™ Cells (ThermoFisher #C640003). The plasmid DNA was sequenced to confirm library and barcode representation and distribution.

##### Spliceosome library annotation

CRISPR-Cas9 base editing outcomes were predicted according to the following rationale. We assumed that if editing occurs for a given sgRNA, all cytosines within the editing window (position 3-8) will be mutated to thymine, with the exception of Cs at positions 3, 4, 6, 7, and 8 if they are preceded by a G(Kluesner et al., 2018). This was used to classify sgRNAs into non-editing (= sgRNAs containing no C within editing window), non-editing_GC (= sgRNAs containing C’s in GC context unfavorable to editing, not at position 5 within sgRNA), and editing sgRNAs. Editing sgRNAs were further classified by MNV (multiple nucleotide variant) prediction using VEP (McLaren et al., 2016). For each MNV, where available only the outcome for MANE (Matched Annotation between NCBI and EBI) transcript was considered. In a next step, consequences were binned into categories and consequence severity was given in this order: CDS_missense > stop_gained > start_lost > SS_acceptor, SS_donor, SS_region, CDS_silent > 3’UTR > 5’UTR. It should be noted, that VEP considers a splice site region variant a sequence variant with a mutation within 1-3 bases of the exon or 3-8 bases of the intron.

##### Virus production and MOI determination

For lentivirus generation and packaging, media for HEK293T cells was supplemented with non-essential amino acids (Gibco #25300054). Cells were seeded 24 h before transfection with jetPRIME reagent (Polyplus #114-15) at a 2.5 μl to 1 μg DNA ratio. Media was changed 6 h post transfection and fresh media was supplemented with ViralBoost Reagent (Alstem #VC100).

The packaging mix consisting of psPAX2 (Addgene #12260) and pMD2.G (Addgene #12259) was prepared at a molar ratio of 1:1. The following reagents were adapted according to scale of lentivirus production:

24-well plate: 2e5 cells seeded, 450 ng target DNA + 450 ng packaging mix per well. 6-well plate: 7.5e5 cells seeded, 1 μg target DNA + 1 μg packaging mix per well. 10 cm plate: 5.2e6 cells seeded, 5.5 μg target DNA + 4.5 μg packaging mix per dish.

Virus was generally concentrated using Lentivirus Precipitation Solution (Alstem #VC100). Virus was titered by seeding 2e5 eHAP FNLS cells in 1 ml media per 6-well plate and immediately adding sequentially diluted virus amounts. 48 h post-transduction the number of BFP positive cells was assessed by flow cytometry. A viral does resulting in 30-40% transduction efficiency, corresponding in an MOI of ~0.3, was used for all subsequent experiments.

##### Generation of eHAP FNLS cell line

Lentivirus was generated with pFNLS and transduced on eHAP cells. Cell lines were selected with blasticidin four days post-transduction for one week and then single cell sorted to obtain monoclonal cell lines. Editing rate of a cell line was assessed by transduction with control sgRNAs (EMX1, HEK2, HEK3, HEK4) and evaluation using sanger sequencing of the editing window and EditR (Kluesner et al., 2018). Clonal cell lines showing high rates of base editing were treated with 10 μM 10-deacetylbaccatin-III (Selleckchem #S2409) for 10-15 days with ploidy assessed every second day. Treatment was stopped as soon as an exclusively haploid cell population was achieved. Editing rate was re-assessed and no changes were observed.

##### Generation of eHAP FNLS mutant cell lines

eHAP FNLS cells were transduced with lentivirus carrying a single sgRNA at an MOI of 0.3. After 2 days sgRNA carrying cells were selected using puromycin for four days. Cells were given two days to recover from selection pressure and then seeded as single cells by limited dilution. The SF3B1 T1080I and PHF5A 2xTL mutations were obtained by first treating cells for 2 weeks with 2 nM PB to enrich for the mutations.

Single cell colonies were maintained and expanded while assessing ploidy and genotyping the clones. Genotyping was performed by using QuickExtract™ DNA Extraction Solution (Lucigen #QE09050) on a fraction of a clone. The region of interest was amplified using custom primers and sanger sequenced. After expansion of the single cell, ploidy was again assessed before freezing the cell line for long term storage.

##### Ploidy assessment for eHAP cell lines

After harvesting, cells are washed with flow cytometry buffer (1x DPBS with 2% FBS, 4 mM EDTA pH 8) and stained on ice with 0.1% sodium citrate, 0.1% Triton X-100, 50 μg/ml propidium iodide for 5 min and immediately assessed by flow cytometry(Beigl et al., 2020).

##### Spliceosome-wide CRISPR-Cas9 base editing screen

The screen was performed at 500x sgRNA representation for entire duration. 66 million eHAP FNLS cells were infected with the lentiviral spliceosome library (marked with BFP) at an MOI of ~0.3, such that every sgRNA was represented in approximately 500 cells. Puromycin selection was started at 48 h post transduction. At six days post-transduction, cells were assessed by flow cytometry to only contain sgRNA carrying cells (BFP-positive cells >95%). Cells were then split into treatment arms (DMSO vs. 2 nM PB; with identical DMSO concentration in both treatment arms). Cells were propagated and treatment was renewed every second day.

At screen end point cells were harvested and gDNA was extracted with QIAamp DNA Blood Maxi Kit (Qiagen #51194). Genome weight was estimated based on measured ploidy of cells. The sequencing library was prepared using NEBnext^®^ Ultra II Q5 Master Mix (NEB #M0544) and custom primers (forward: IB0096-IB0104; reverse IB0106-IB0121) to have a balanced read sample. A barcode in the reverse primer was used for identification of the sequencing libraries. Libraries were gel-purified and cleaned up. Libraries were balanced and quality was assessed with Bioanalyzer High Sensitivity DNA Kit, Agilent #5067-4626).

##### Validation experiments

For validation experiment, 45 individual sgRNAs targeting the spliceosome were cloned into pLibrary containing EF1α-puro-T2A-BFP and made into lentivirus as described above. sgRNAs were selected for significant enrichment or depletion (LFC > |2|, padj < 0.05). Moreover, we took statistically not significantly enriched/depleted sgRNAs if they were strongly enriched (LFC > 2.75) or depleted (LFC < −3.5). The threshold was set to approximately mimic the lowest level of enrichment or depletion observed, respectively, for the statistically significant sgRNA. All depleted sgRNAs further had to fulfil LFC > −1 for a comparison of t14 vs. t0 in the control condition. In addition, three non-targeting sgRNAs were cloned into pLibrary containing EF1α-puro-T2A-mCherry and into pLibrary EF1α-puro-T2A-BFP (see Table S3 for full list and primers). Lentivirus was generated and titered as described above. 2e5 eHAP FNLS cells were transduced with individual sgRNA lentivirus in 6-well plates. Puromycin selection was started 36 h post-transduction and continued for four days. Care was taken to maintain all cells haploid (as diploid cells grow faster) and ploidy was checked with flowcytometry as described above. On day 6 (= t0) cells were grouped according to their growth density on plate and a representative sample was counted. An estimated 5500 cells per spliceosome sgRNA or non-targeting sgRNA carrying cells were each mixed with 5500 cells carrying sgNTC_400.

Treatment was started on the next day (DMSO vs. 2 nM PB) and cells were passaged as needed with treatment renewed every second day. The ratio of BFP:mCherry was assessed at t0, t4, t8 and t15.

##### Library and sequencing for base editing window

Lentivirus from validation experiment was used to transduce 2e5 eHAP FNLS cells at MOI 0.3 in a 6-well dish. After puromycin selection on day 6 (= t0), 2x 11000 cells were seeded in 96-well plates and split into treatment arms the next day (DMSO vs. 2 nM PB). Cells were propagated and harvested at t8 and t14 for gDNA extraction. Genomic DNA was extracted using QuickExtract™ DNA Extraction Solution (Lucigen #QE09050) and 1 μl (qsp. >200 cells) and was used for target site amplification using a 2-step PCR. In addition, each reaction contained 10 μl NEBnext Ultra II Q5 master mix as well as 1 μM of each forward and reverse primer. Primer pairs for PCR 1 were selected such that forward annealing primer is not closer than 7 nt to editing window position 1 but still allowing that 75 sequencing cycles will read the sequence. Primers were verified to anneal to a single position within the genome using BLAT (Kent, 2002) and tested before use. Primers for PCR 1 (12 cycles) were flanked for the forward primer by: 5’-ACACTCTTTCCCTACACGACGCTCTTCCGATCT-[target specific sequence]-3’ and for the reverse primer by 5’-GTGACTGGAGTTCAGACGTGTGCTCTTCCGATCT-[target specific sequence]-3’. 1.5 μl of PCR 1 were used as template for PCR 2 (14 cycles) and used forward primers 5’-AATGATACGGCGACCACCGAGATCTACAC-NNNNNNNN-ACACTCTTTCCCTACACGAC-3’ (compatible to Illumina i5) and reverse primers 5’-CAAGCAGAAGACGGCATACGAGAT-NNNNNNNN-GTGACTGGAGTTCAGACGTG-3’ (compatible to Illumina i7), with both containing 8N barcodes for multiplexing.

Primers were removed after the second PCR step with AMPure XP Reagent (Beckman #A63882). Library was quantified (QuantiFluor^®^ dsDNA System; Promega #E2670) before being pooled (10 fmol per sample) for each time point & condition, run on 8% TBE-PAGE gel, then size selected for a range of 250-500 bp. Pooled libraries were then run on Bioanalyzer High Sensitivity DNA Kit, Agilent #5067-4626) before final pooling and subsequent run on a MiniSeq Sequencing System using the Miniseq High Output Kit (75 cycles) (Illumina #FC-420-1001) with a 10% of phiX spike-in.

##### RNA-seq

eHAP FNLS and mutant cells were treated for 3 h with DMSO or 2 nM PB before cells were harvested. This was done for three replicates of an eHAP FNLS cell line transduced with a non-targeting sgRNA. Total RNA was extracted using the RNAqueous^TM^-96 Total RNA Isolation Kit (ThermoFisher #AM1920) according to the manufacturer’s protocol. Poly(A)-enriched RNA was obtained with the poly(A) RNA Selection Kit V1.5 (Lexogen #157.96) and RNA-seq libraries were generated using the CORALL Total RNA-Seq Library Prep Kit (Lexogen #117.96). The libraries were then sequenced using paired-end 150 bp reads with 60 million reads per sample on Nova Seq (S4).

##### RNA extraction and RT-PCR

RNA was extracted from cells using the RNeasy Plus Mini Kit (Qiagen #74134) according to the manufacturer’s protocol. Reverse transcription using either SuperScript III (Invitrogen #18080093) or SuperScript IV (Invitrogen #18090050) was performed according to the manufacturer’s protocol using a mix of random hexamer primers and oligo(dT) using an input of 500 ng total RNA for a 10 μl reaction. PCR was performed with junction specific primers (Table S3) using 2% of the cDNA as an input for a 25 μl PCR reaction. PAGE was visualized using SYBR Gold (Invitrogen #S11494) and intensity of PCR products were quantified using ImageJ (NIH).

##### BioID sample preparation for MS

6 million HEK293T cells were seeded on 150 mm plates and transfected 24 hours later with 15 μg plasmid using jetPRIME reagent (at a 1:2.5 ratio for μg DNA:μl jetPRIME reagent; Polyplus #114-15). Transfection was performed in four independent biological replicates. Media was replaced with fresh culture media 5-6 h post transfection. At 24 h post transfection, a biotin pulse of 2 min biotin (culture media supplemented with 200 μM biotin) was used for proximity labelling. Cells were then immediately placed on ice and washed five times with ice cold DPBS (Corning #21-031-CV) before collection by gentle repeat pipetting. Cell pellets were lysed in 1500 μl ice cold RIPA buffer (50 mM Tris-Cl pH 7.4, 150 mM NaCl, 1% NP40, 0.5% Na-deoxycholate, 0.1% SDS, 1 mM EDTA, Roche cOmplete and 1 mM PMSF). Cell lysates were clarified by centrifugation (13000 x g at 4°C for 10 min) before quantification with BCA protein assay (Thermo #23227). 2.5 mg protein in 1500 μl RIPA buffer were added to 250 μl MyOne Streptavidin T1 Dynabeads (Invitrogen #65602), prewashed twice with 1 ml RIPA buffer. Protein and beads were incubated for 30 min at 4 °C with rotation to capture biotinylated proteins. Beads were then pelleted on magnet and washed twice with RIPA buffer (1 ml for 2 min at RT), washed once with 1 M KCL (1 ml for 2 min at RT), washed once with freshly made 0.1 M Na_2_CO_3_ (1 ml for 10 s), washed once with freshly made 2 M urea in 10 mM Tris.Cl pH 8 (1 ml for 10 s), before washing twice more with RIPA buffer (1 ml per wash, 2 min at RT). Beads were resuspended in 200 μl RIPA buffer before transfer to a new low binding tube. Beads were then washed in 200 μl 50 mM Tris.Cl pH 7.5, twice in 200 μl 2 M urea in 10 mM Tris.Cl pH 7.5, and twice in 200 μl H_2_O.

Bead pellet was frozen for handover to MS facility.

##### On beads Digestion and TMT labelling

Sample-incubated streptavidin magnetic beads were resuspended in 9 μl 5 mM Tris(2-carboxyethyl)phosphine 20mM triethylammonium bicarbonate and incubated for 30 min at room temperature. After this, iodoacetamide was added to a final concentration of 7.5 mM, and samples incubated for 30 additional minutes. 1 μg of LysC (Fujifilm Wako Pure Chemical Corporation) was added to each sample and incubated at 37 °C overnight. Then 1 μg sequencing grade trypsin (Promega) was added to each sample and incubated at 37 °C overnight. Supernatants of the beads were recovered, and beads digested again using 0.5 ug trypsin in 100mM NH_4_HCO_3_ for 2 h. Peptides from both consecutive digestions were combined and recovered by solid phase extraction using C18 ZipTips (Millipore), eluted in 15 μl 50% acetonitrile 0.1% formic acid, and evaporated. Samples were then resuspended in 8 μl 0.1 M triethylammonium bicarbonate pH 8.0. Dried samples were labelled according to TMTPro^TM^-16 label plex kit instructions (ThermoFisher Scientific). Briefly, TMT reagents were dissolved in acetonitrile at 12.5 μg/μl, and 4 μl of these stocks added to the samples. After incubation for 1 h at room temperature samples were quenched with 1ul 5% hydroxylamine, and all 16 samples were combined, partially evaporated, and desalted using a C18 ZipTip as described before. The eluate was dried in preparation for LC-MSMS analysis.

##### Mass Spectrometry Analysis

Samples coming from RP fractionation were run onto a 2 μm, 75μm ID x 50 cm PepMap RSLC C18 EasySpray column (Thermo Scientific). 3-hour MeCN gradients (2–30% in 0.1% formic acid) were used to separate peptides, at a flow rate of 300 nl/min, for analysis in a Orbitrap Lumos Fusion (Thermo Scientific) in positive ion mode. MS spectra were acquired between 375 and 1500 m/z with a resolution of 120000. For each MS spectrum, multiply charged ions over the selected threshold (2E4) were selected for MSMS in cycles of 3 seconds with an isolation window of 0.7 m/z. Precursor ions were fragmented by HCD using stepped relative collision energies of 30, 35 and 40 to ensure efficient generation of sequence ions as well as TMT reporter ions. MSMS spectra were acquired in centroid mode with resolution 60000 from m/z=110. A dynamic exclusion window was applied which prevented the same m/z (mass tolerance 30 ppm) from being selected for 3 0s after its acquisition.

##### Immunoblotting

Cell lysates were mixed with NuPAGE LDS Sample Buffer (Invitrogen #NP00007), heated for 5 min at 95 °C, separated on SDS-PAGE gels and transferred to nitrocellulose membranes. Blots were either incubated for 1 h at RT or overnight at 4 °C with the following primary antibodies in TBS-T with 5% milk: α-HA (Cell Signalling Technology, #3724) at 1:5000; α-GAPDH-HRP (Proteintech, #HRP-60004) at 1:10,000. HRP-conjugated streptavidin (Invitrogen, #S911) reconstituted at 1 mg/ml was used at 0.3 μg/ml in 3% (w/v) BSA in 1x TBST and blots were only incubated for 30 min in its presence.

##### Protein expression and purification

For pulldown and RNA binding assays, MBP-*hs*SUGP1 variants were expressed *E. coli* BL21 Star (DE3) (Invitrogen). Cells were grown at 37°C in LB medium until an OD_600_ of 0.6 was reached. Protein expression was induced with 2□mM isopropyl-ß-D-thiogalactopyranoside (IPTG) and maintained at 37°C for 3 h. Expression cultures were harvested by centrifugation and cell pellets were resuspended in lysis buffer (50 mM Hepes, pH 7.5, 200 mM NaCl, 2 mM dithiothreitol [DTT]) supplemented with cOmplete EDTA-free protease inhibitor mixture (Roche), 1 mg/mL lysozyme (Sigma), and 5 μg/mL DNaseI (Roche). For cell lysis, the suspension was passaged through a LM10 Microfluidizer. Subsequently, the lysate was cleared by centrifugation at 3200 g for 10 min and filtered (0.45 μm). Lysates were incubated with preequilibrated amylose beads (New England BioLabs) for 1 h at 4°C. Beads were washed with lysis buffer and bound proteins were eluted with lysis buffer containing 25 mM maltose. The eluates were concentrated, loaded onto a gel-filtration column (Superdex 200, GE Healthcare) and eluted in size-exclusion buffer (10 mM Hepes, pH 7.5, 200 mM NaCl, 2 mM DTT). The MBP-*hs*SUGP1 variants were either directly used in biochemical assays or flash-frozen in liquid nitrogen and stored at −80°C.

For the ATPase assay, MBP-*hs*SUGP1 variants were subjected to additional washes while bound to the amylose beads during the first purification step to remove ATPase contamination. Bound proteins where incubated with lysis buffer supplemented with 2 mM ATP and 2 mM MgCl_2_ for 10 min at 4°C and washed with lysis buffer containing 1 M NaCl. After the high-salt wash, beads were equilibrated again in lysis buffer before elution as described above.

##### Protein binding assays

For interaction studies, purified His_10_-*hs*DHX15ΔN and MBP-*hs*SUGP1 variants were mixed in equimolar amounts in pulldown buffer (50 mM Hepes, pH 7.5, 200 mM NaCl, 2 mM DTT). His_6_-MBP was used as a negative control. Proteins were incubated with 50 μl of amylose beads (50% slurry in pulldown buffer) for 1 h at 4°C on a rotator. The beads were washed three times in pulldown buffer. Bound proteins were eluted with pulldown buffer containing 25 mM maltose. Proteins of input and eluate samples were separated by sodium dodecyl sulfate (SDS)/polyacrylamide gel electrophoresis (PAGE) and were visualized by Coomassie staining.

##### RNA Binding Assays

RNA binding affinities of His_10_-*hs*DHX15ΔN in complex with MBP-*hs*SUGP1 variants were determined by measuring changes of fluorescence polarization (FP) in dependence of protein concentration, as previously described (Studer et al., 2020b). Experiments were performed in binding buffer (20 mM Hepes (pH 7.5), 150 mM NaCl, 5% glycerol and 2 mM MgCl_2_) with 10 nM 5’-6-fluorescein amidites (FAM)–labelled U_12_ RNA (Microsynth) and protein concentrations ranging from 1 nM to 32 μM. FP was determined using a CLARIOstar microplate reader (BMG Labtech) by excitation at 482 nm and detection at 530 nm wavelength. All samples were measured five times and all samples were prepared in triplicates. After baseline subtraction, the obtained FP values were normalized to 1 and fitted according to Rossi *et al.* using GraphPrad Prism (Rossi and Taylor, 2011).

##### ATPase assays

Activity of *hs*DHX15 and its interactor *hs*SUGP1 were monitored using an NADH-coupled ATPase assay. For all measurements of ATPase activity, 1.8 μM His_10_-*hs*DHX15ΔN and 1.8 μM MBP-*hs*SUGP1 variants (G-patch (548-611) were mixed in ATPase buffer containing 50 mM Hepes pH 7.5, 50 mM KAc, 5 mM MgAc_2_, 2 mM DTT, 0.5 mM nicotinamide adenine dinucleotide, 1 mM phosphoenolpyruvate, 12 U of pyruvate kinase, and 18 U of lactate dehydrogenase. All measurements were carried out in half area 96-well plates (Greiner). After equilibration for 10 min at 37°C, reactions were started by adding 250 μM ATP (pH 7.5). Absorption at 340 nm was measured over a time course of 40 min at 37°C with one measurement per minute using a CLARIOstar microplate reader. Absorption change in the absence of ATP was measured for baseline correction. Absorption values were adjusted to a path length of 1 cm. Absorption change over time was determined by linear regression and converted to concentration change over time with an extinction coefficient at 340 nm of 6,220 M^-1^cm^-1^ using Beer-Lambert’s law. Initial velocities were derived from concentration change over time using a total enzyme concentration of 1.8 μM. All measurements were prepared in triplicates. Absence of ATPase contaminations from *hs*SUGP1 preparations was confirmed by measuring ATPase activity in the absence of *hs*DHX15ΔN.

#### DATA ANALYSIS

##### Screen analysis

The sequencing data was demultiplexed to obtain individual samples for timepoints and trimmed (BBMap BBDuk v.38.94). Reads were counted by alignment to a reference file of all sgRNAs present in the pool. In the next step, each barcode was randomly assigned to either of two replicates. 32 sgRNA targeted 2 loci and their reads were duplicated and assigned to both targets. Log2-fold changes between samples were calculated using DESeq, filtering out reads with on average less than 50 reads across all samples(Love et al., 2014).

##### Identification of causative mutations

Editing at the base editing window was quantified using CRISPResso2 v.2.1 (Pinello et al., 2016), run with the following parameters: [--exclude_bp_from_left 18 --exclude_bp_from_right 0 --quantification_window_center −12 --quantification_window_size 15 --min_average_read_quality 30 --default_min_aln_score 60 --plot_window_size 20 --base_editor_output --output_folder 20210531_pos -p 12]. For analysis we used the “Alleles_frequency_table_around_[sgRNA].txt” output files or in the single case where editing occurred far outside the editing window “Alleles_frequency_table.txt” was used for information extraction.

Samples were processed to only consider those with 100 or more reads per condition across all conditions. In addition, all alleles with <1% in all conditions were removed before re-calculating percent distribution of alleles. Alleles were translated and grouped by protein sequence outcome and for each percent distribution was summed. Log2-fold change was calculated for samples

###### Validation of PB sensitive samples

For samples depleted in the screen, only sequencing data corresponding to t0 were considered. Mutational outcome was noted for any event with >5% frequency.

###### Validation of PB resistant samples

All timepoints and treatment conditions were considered for the analysis.

##### Splicing analysis

The RNA-seq data was demultiplexed, trimmed (BBMap BBDuk v.38.94), and then mapped using STAR 2.7.9 (Dobin et al., 2013) [--alignSJoverhangMin 8 --alignSJDBoverhangMin 1 --outFilterMismatchNmax 999 --alignIntronMin 20 --alignIntronMax 1000000 --alignMatesGapMax 1000000 --peOverlapNbasesMin 10 --peOverlapMMp 0.2] and SAMtools (v1.10) for conversion to sorted bam files. Reads were then deduplicated using the UMIs from the Lexogen CORALL workflow to prevent removal of natural duplicates. On average, 85% of the reads could be mapped and quality assessment using the RSeQC package (Wang et al., 2012) showed neither a bias in read distribution nor gene body coverage. Differential expression of genes was assessed using RSubreads (Liao et al., 2019) for counting and DESeq2 (Love et al., 2014). To analyze splicing at the exon level only junctions with more than 10 junction reads were considered. We used rMATS-turbo (Shen et al., 2014a) (hereafter, rMATS) for quantification. For all analysis performed with rMATS only splicing junctions with FDR < 0.01 were considered. Alternatively, we also used MAJIQ(v2.4) (Mehmood et al., 2020) with |dPSI] ≥10 and Confidence Threshold > 90%. For analysis of intron and exon features we used the software matt (Gohr and Irimia, 2019). When preparing figures with IGV, junction reads with low numbers were removed for clarity

##### Peptide and protein identification and TMT quantitation

Peak lists were generated using PAVA in-house software (Guan et al., 2022). All generated peak lists were searched against the human subset of the SwissProt database (SwissProt.2019.07.31), using Protein Prospector (Clauser et al., 1999) with the following parameters: Enzyme specificity was set as Trypsin, and up to 2 missed cleavages per peptide were allowed. Carbamidomethylation of cysteine residues, and TMTPro16plex labelling of lysine residues and N-terminus of the protein were allowed as fixed modifications. N-acetylation of the N-terminus of the protein, loss of protein N-terminal methionine, pyroglutamate formation from of peptide N-terminal glutamines, and oxidation of methionine were allowed as variable modifications. Mass tolerance was 10 ppm in MS and 30 ppm in MS/MS. The false positive rate was estimated by searching the data using a concatenated database which contains the original SwissProt database, as well as a version of each original entry where the sequence has been randomized. A 1% FDR was permitted at the protein and peptide level. For quantitation only unique peptides were considered; peptides common to several proteins were not used for quantitative analysis. Relative quantization of peptide abundance was performed via calculation of the intensity of reporter ions corresponding to the different TMT labels, present in MS/MS spectra. Intensities were determined by Protein Prospector. Summed intensity per sample on each TMT channel for all identified carboxylases were used to normalize individual intensity values. Relative abundances were calculated as ratios vs the average intensity levels in the 4 channels corresponding to control samples. For total protein relative levels, peptide ratios were aggregated to the protein levels using median values of the log2 ratios.

##### Software for data visualization and statistical analyses

For visualisation of cryo-EM and crystal structures we used PyMOL (v2.3.5), for visual presentation we used R ggplot2 (v3.3.6) with MetBrewer (v0.2.0.) and ggVennDiagram (v1.2.0) or pheatmap (v1.0.12), and GraphPad Prism (v8), and the Integrative Genomics Viewer (IGV) (v2.12.2). Schematics were created with Adobe Illustrator 2023.

All statistical analyses were performed in R (v4.1.2) using base R or package multcomp. Statistical significance of one-way ANOVA or t-test is indicated by asterisks (* p < 0.05, ** p < 0.01, *** p < 0.001), unless otherwise indicated.

## SUPPLEMENTARY FIGURE LEGENDS

<u>Figure S1. Spliceosome base editor screen</u>

(A) Lentiviral vectors used in this study. *Left:* Construct for generation of eHAP FNLS cell line by transduction and selection for blasticidin resistant cells exhibiting GFP fluorescence. *Right:* Vector for pooled library. The sgRNA is followed by a N_20_-barcode which will uniquely label each sgRNA and can be read out during paired-end sequencing. The vector carries a puromycin resistance cassette for easy selection during the screen.

(B) Schematic of CRISPR-Cas9 base editing. The sgRNA loaded on a nickase Cas9 (nCas9) targets a tethered ssDNA-specific deaminase to a chosen genomic locus. The deaminase then modifies all nucleotides within the editing window.

(C) Benchmark test of eHAP1 FNLS cell line for four commonly used sgRNAs. Shown is the rate for C > T editing as well as for C > R transversion mutations six days post-transduction with the sgRNA and after selection for sgRNA carrying cells. Overall high rates of editing are observed for eHAP FNLS. Editing rates across replicates are very consistent. (n = 4, except HEK2 n=6)

(D) Benchmark test of eHAP1 FNLS cell line. Shown are Sanger sequencing traces for the target region of EMX1 and HEK3 sgRNA. In the presence of the sgRNA the cytosines within the editing window (position 5 and 6, or 4 and 5 respectively) are modified.

(E) Fraction of nucleotides that can be modified across the coding sequence targeted by the spliceosome sgRNA library with FNLS.

(F) Predicted editing consequences per sgRNA from VEP when utilizing FNLS. ~70% of changes are predicted to be of high impact. Consequences were binned into categories based on most severe outcome with splice site acceptor/donor variant > nonsense > stop lost > start lost > missense > splice region variant > silent > intron variant > UTR variant.

(G) Performance of sgRNAs targeting spliceosomal proteins, binned by predicted editing consequence. Non-targeting sgRNAs serve as a negative control. The dashed line marks the LFC for the bottom 5% of negative controls (LFC = −0.8814), percentage of sgRNAs falling below this threshold is indicated. Figure drawn in R with ggplot2 (v3.3.6) and ggridges (v0.5.3).

<u>Figure S2 Validation and mapping experiments</u>

(A) Indivdual sgRNAs and their performance in the confirmation assay. Measurements are from three independent transductions (n=3). Student’s t-Test (paired) with *P* value * p < 0.05, ** p < 0.01, and *** p < 0.001, respectively.

(B) Sum of conversion per each position within the protospacer, for 23 sgRNAs validated for hypersensitivity to PB. The position of the C is indicated along the x axis, with 1 corresponding to the first nucleotide of the protospacer and 21-23 corresponding to the PAM. The editing window (3-8) is shown in gray. Lines indicate median of edits at that position across all 23 sgRNAs.

(C) Location of PB sensitive mutations in SF1 mapped to structure of chimeric SF1 SURP binding region fused to SF3A1 SURP1. SF1 residues S315 and L316 are targeted by sgSF1_265, which gives rise to PB sensitivity. The target lies within the SF1 SURP binding region and forms part of the hydrophobic interface with SF3A1 SURP1. sgSF3A1_121 and sgSF3A1_283 result in mutation of SF3A1 SURP1 residues S110 and R50, respectively, which results in PB sensitivity. All sgRNA targeted residues are shown in magenta. (PDB: 7VH9)

(D) Location of the SURP1 domain of SF3A1 in the B^act^ complex, the earliest stage of spliceosome assembly it has been visualized in. PB hypersensitive mutations of SF3A1 target R50 and S110, which locate to the SURP1 domain. Both residues are at this stage in proximity to XAB2/SYF1 and CRINKL/SYF3 which are part of “bridge 2” as described by Haselbach *et al.* (2018). (PDB: 6FF7)

(E+F) Six sgRNAs targeting CDC40/hPrp17 give rise to PB sensitivity. (PDB: 5XJC) (E) Three target the N-terminal domain of CDC40 at the flexible N-terminus (S14 and S16) as well as at P95 and G113. P95 is in close contact to the PPIL1 domain, while G113 is at the interface to RBM22. (F) Three more sgRNAs target the C-terminal WD40 domain of CDC40. The target sites are located in loops which face away from the catalytic core.

<u>Figure S3. Analysis of mutations in PHF5A</u>

(A) Editing outcome for sgPHF5A_7: In the absence of PB the sgRNA results in a K29N mutation. PB treatment further enriches for this mutation. Bystander mutations at D27 may occur but do not strongly enrich.

(B) Editing outcome for sgPHF5A_21: In the absence of PB some mutations occur at E74. PB treatment enriches for an E74D mutation (with E74N also showing some enrichment vs. the DMSO control sample).

(C) Editing outcome for sgPHF5A_47: In the absence of PB the 3’SS of exon 3 is mutated. PB treatment enriches for mutations at D27.

(D) Editing outcome for sgPHF5A_26: In the absence of PB mutations occur at S67. PB treatment enriches for an S67C which is present at a similar prevalence as S67F at t15.

(E) Comparison of predicted mutations in PHF5A with actual confirmed mutations in PHF5A. Deep sequencing of the target region allows for the precise identification of mutations likely responsible for phenotype (PB resistance). This shows that while sgRNAs can predict the region where mutations occur, in this instance only D27 could be confirmed as a mutation site. The four other sgRNAs resulted in mutations proximal to the predicted editing sites.

<u>Figure S4. Clonal cell line generation and analysis</u>

(A) Schematic for generation of monoclonal eHAP FNLS cell lines carrying mutations in splicing factors. All cell lines were genotyped and verified to be haploid.

(B) Sixty hour cell proliferation profiling (CellTiter-AQueous cellular viability and cytotoxicity assay) of control eHAP FNLS cell line expressing non-targeting sgRNA and monoclonal cell lines carrying either SUGP1 E554K or SUGP1 G603N mutation to PB. Error bars indicate s.d. n = 3 (average of two technical replicates for independent clonal cell lines).

(C) Cell cycle analysis for eHAP FNLS upon treatment with 2 nM PB. After 4 h an increase in G2% suggests cell cycle arrest and death. Cell cycle analysis was performed on ethanol fixated cells by staining with propidium iodide after RNase treatment (5 μg RNase A at RT for 30 min per max. 1e6 cells).

(D) MA-plots for RNA-seq comparing control cell line transcript mean counts vs. mutant cell lines under DMSO treatment. Mutations do not appear to affect the transcriptome.

(E) Sashimi plot for alternative 3’ splice site usage in *ENOSF1* exon 11 (DMSO). The control cell line exclusively uses the distal 3’SS while the SUGP1 mutants also make use of a proximal 3’SS. Representative traces are shown (plot generated with IGV with only junctions ≥5 reads shown; alignment to Hg38) and junctions of interest are emphasized.

(F) RT-PCR and quantification for alternative 3’ splice site usage for *ENOSF1* exon 11 (DMSO). RNA extracted from eHAP FNLS cell lines – same monoclonal cell lines as for RNA-seq.

(G) PAGE as used for quantification in (F and 5G). Shown is also the RT-control (pooled samples for each genetic background), which shows no amplification.

*Statistical analysis for RT-PCR:* one-way ANOVA with Dunnett’s for multiple comparison (two-sided, with control as reference) was performed with R and package multcomp (v.1.4-20); * p < 0.05, ** p < 0.01, and *** p < 0.001; all with n = 3.

<u>Figure S5. SUGP1 mutant splicing analysis</u>

(A) Expanded view of Figure 5C focusing on SUGP1 mutants.

(B) Quantification of differential splicing in eHAP FNLS control or mutant cell lines by RT-PCR corresponding to observations from RNA-seq as shown in Figure 5 and S5D. Shown is the change in cassette exon splicing upon treatment with PB vs. DMSO. *Top:* PB induces increased exon skipping for *RBM5* exon 16 to the same degree in control as well as the SUGP1 mutant cell lines. *Bottom:* PB induces increased exon skipping for *ORC6* exon 5. SUGP1 mutation modulates this phenotype, with the G603N mutant cell lines showing a significant decrease in exon 5 skipping compared to the control cell lines. Statistical significance is only shown for mutant samples vs. control (= reference).

(C) Representative PAGE as used for quantification in (B).

(D) Cartoon representation of SUGP1 and its domain and motif organisation. α-helices identified with AlphaFold2 and in proximity to the G-patch motif are shown in orange (αH4-5, αH6, and αH7). Several cancer mutations locate to these α-helices or their vicinity (grey arrows and residue numbers).

(E) Western blot for overexpression assays using HA-SUGP1 or HA-SUGP1 mutants as shown in (G-H). 1 μg plasmid was transfected into HEK293T cells and RNA or protein samples were extracted 48 h later. Membrane had to be cropped between samples L570E and ΔαH4-5 (non-discussed sample).

(G) RT-PCR and quantification for alternative 3’ splice site usage in *ENOSF1* exon 11 upon SUGP1 wt or mutant protein overexpression (48 h) in HEK293T cells. Overexpression of SUGP1 wt does not influence splicing outcome. αH6 shows an equally strong phenotype as G-patch point mutants. A representative example of the PAGE used for quantification is shown.

(H) RT-PCR and quantification for alternative 3’ splice site usage in *ENOSF1* exon 11 upon SUGP1 wt or mutant protein overexpression (48 h) in HEK293T cells. Overexpression of SUGP1 wt does not influence splicing outcome. All mutants affect splicing 3’ splice site choice, with αH4-5 and αH6 showing a similar phenotype as G603N. A representative example of the PAGE used for quantification is shown.

*Statistical analysis for RT-PCR:* one-way ANOVA with Dunnett’s *(ENOSF1, TMEM14C* – with control as reference) or Tukey’s *(RBM5, ORC6)* for multiple comparison (two-sided) was performed with R and package multcomp (v.1.4-20); * p < 0.05, ** p < 0.01, and *** p < 0.001; all with n = 3.

<u>Figure S6. Proximity labeling experiments</u>

(A) Schematic for BioID experiments.

(B) Optimization of labeling using SUGP1-miniTurboID overexpression in HEK293T cells. Different time points and biotin concentrations were compared to obtain strong labeling within a short time frame. When no SUGP1-miniTurboID was transfected only background levels of endogenously biotinylated proteins are observed.

(C) miniTurbo proximity labeling using SUGP1-miniTurboID L570E vs. wt overexpressed in HEK293T cells. Biotinylated proteins were identified with TMT-MS after enrichment with streptavidin beads. −log_10_(p-value) is plotted against the log_2_-fold change (LFC) in a volcano plot (dashed lines: cutoffs at p < 0.05 and LFC > 0.5). All depleted or enriched proteins are labelled and spliceosome associated factors are underlined. SNRNP48 (associated with the minor spliceosome) and DHX15 are the only spliceosomal proteins that show a significant change, with both depleted. (n = 4)

(D) Domain organization and schematic representation of the MBP-*hs*SUGP1 variants with the introduced mutations colored in red for construct αH6-αH7 (522-633).

(E) Coomassie-stained gels of protein binding assays using purified MBP-*hs*SUGP1 construct and His_10_-*hs*DHX15ΔN. MBP-SUGP1 αH6-αH7 with either wildtype (wt) protein sequence or carrying the indicated mutation were used as baits and His_6_-MBP served as a control. Input (1.5% of total) and eluates (24% of total) were loaded.

(F) Fluorescence polarization of FAM-labeled U_12_ RNA with His_10_-*hs*DHX15ΔN in the absence or presence of MBP-SUGP1 αH6-αH7 wt or mutants. The dashed line indicates 50% normalized polarization. Error bars represent standard deviations from the average values of triplicate measurements. RNA dissociation constants (Kd) with standard error of means (SEM) were derived from fitting the respective data by linear regression.

## Notes

### Competing Interest Statement

The authors have declared no competing interest.

