## Supplementary figures and images for "Targeted high throughput mutagenesis of the human spliceosome reveals its *in vivo* operating principles"

### Supplemental Figure S1

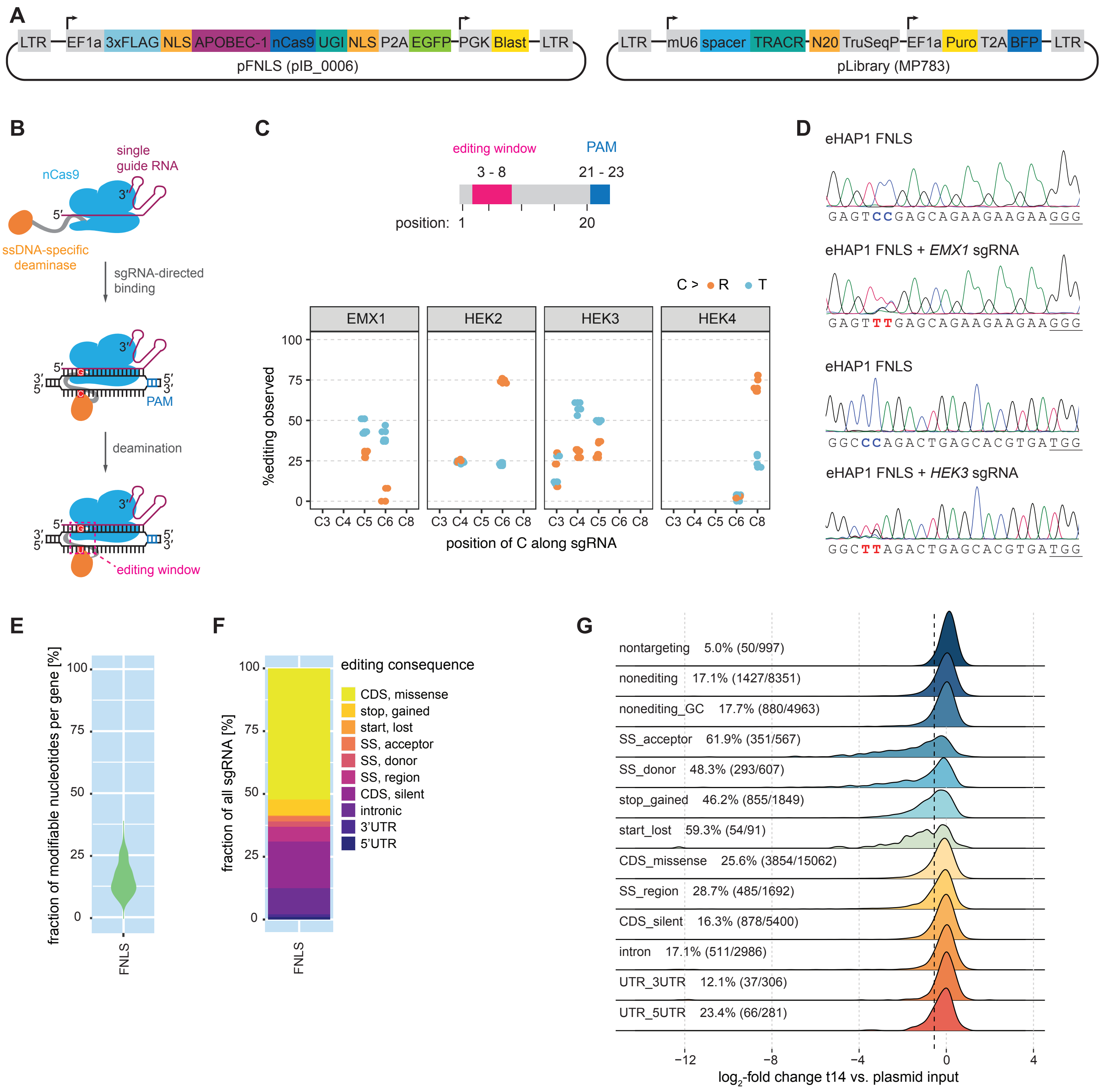

### Supplemental Figure S2

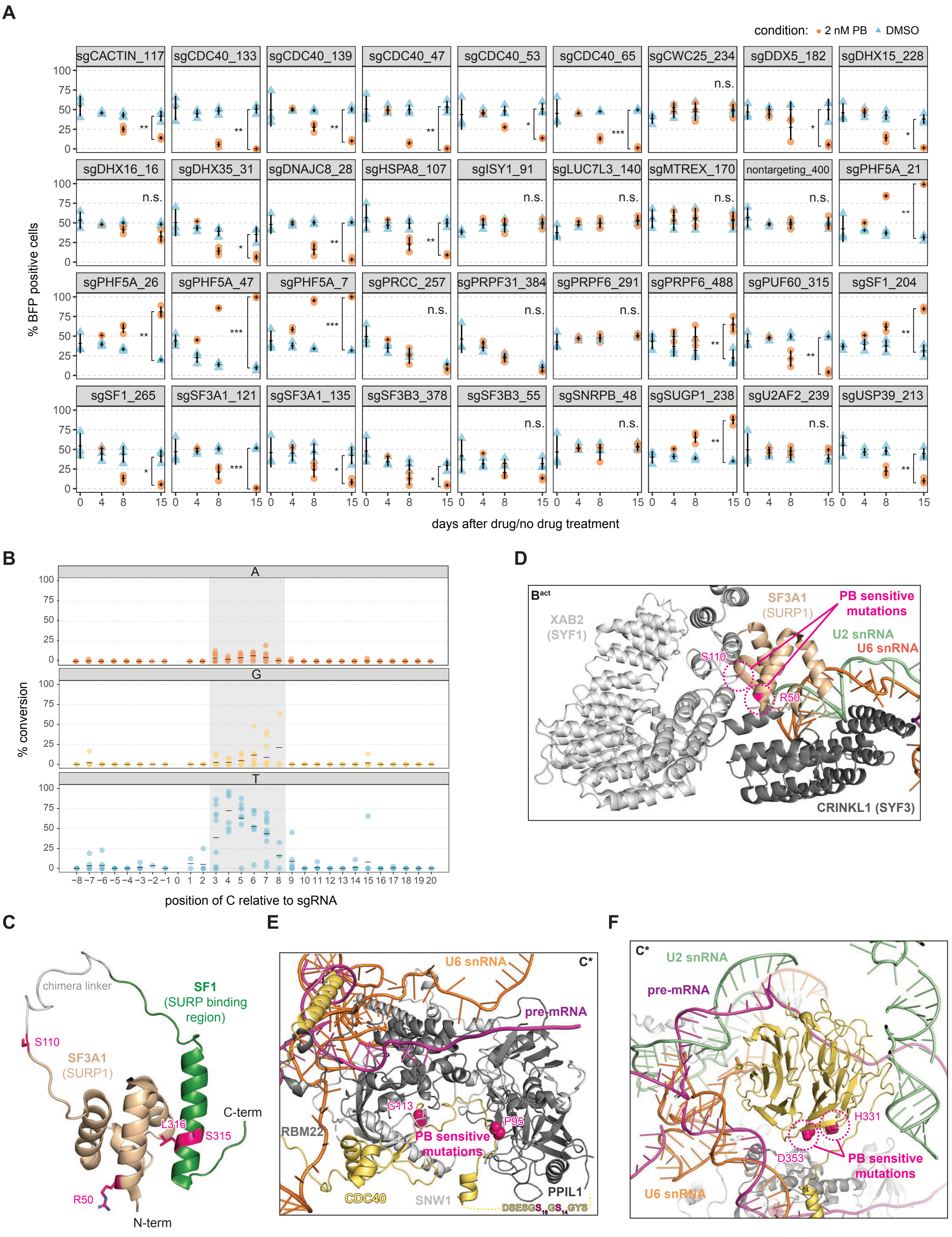

### Supplemental Figure S3

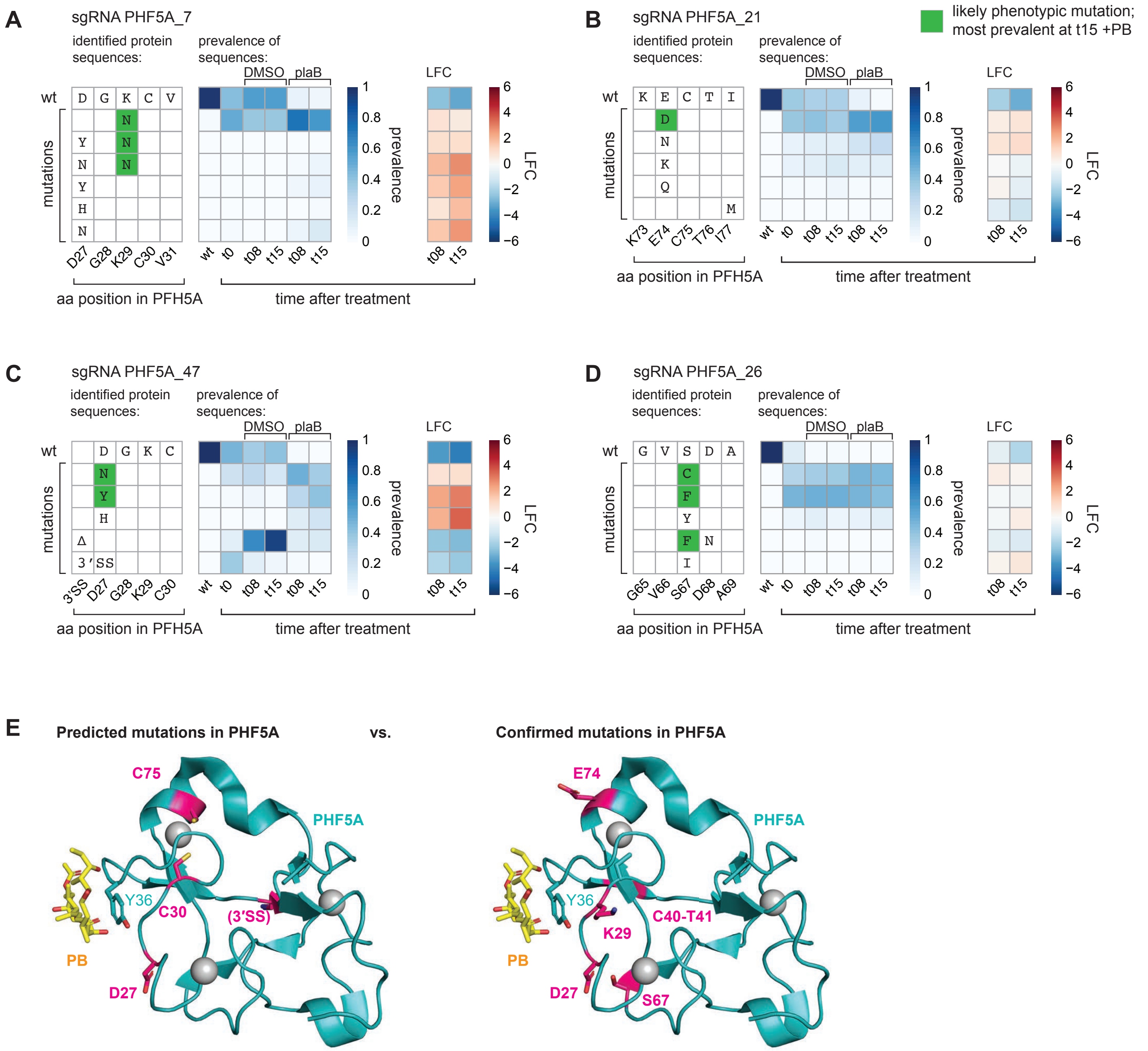

### Supplemental Figure S4

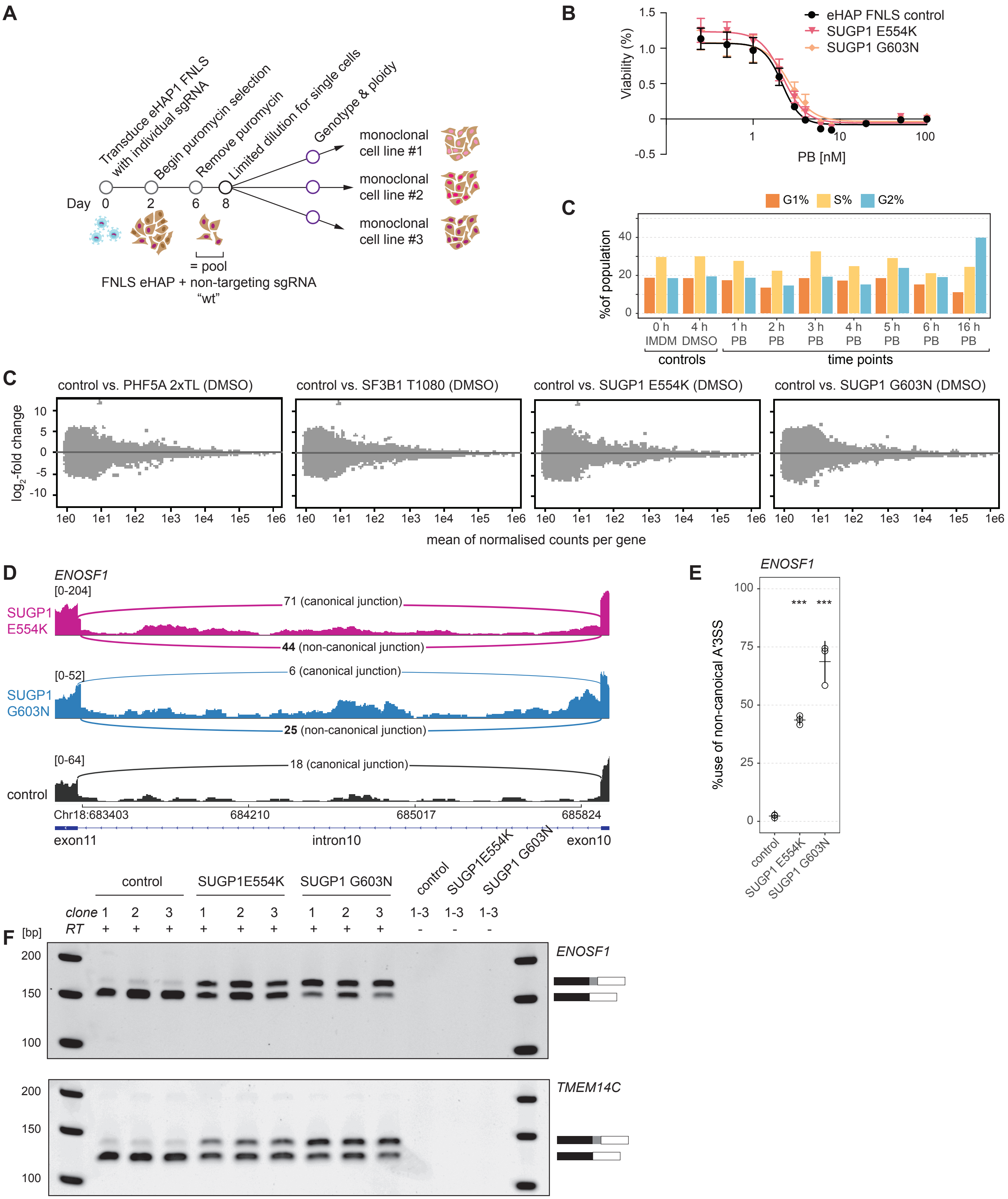

### Supplemental Figure S5

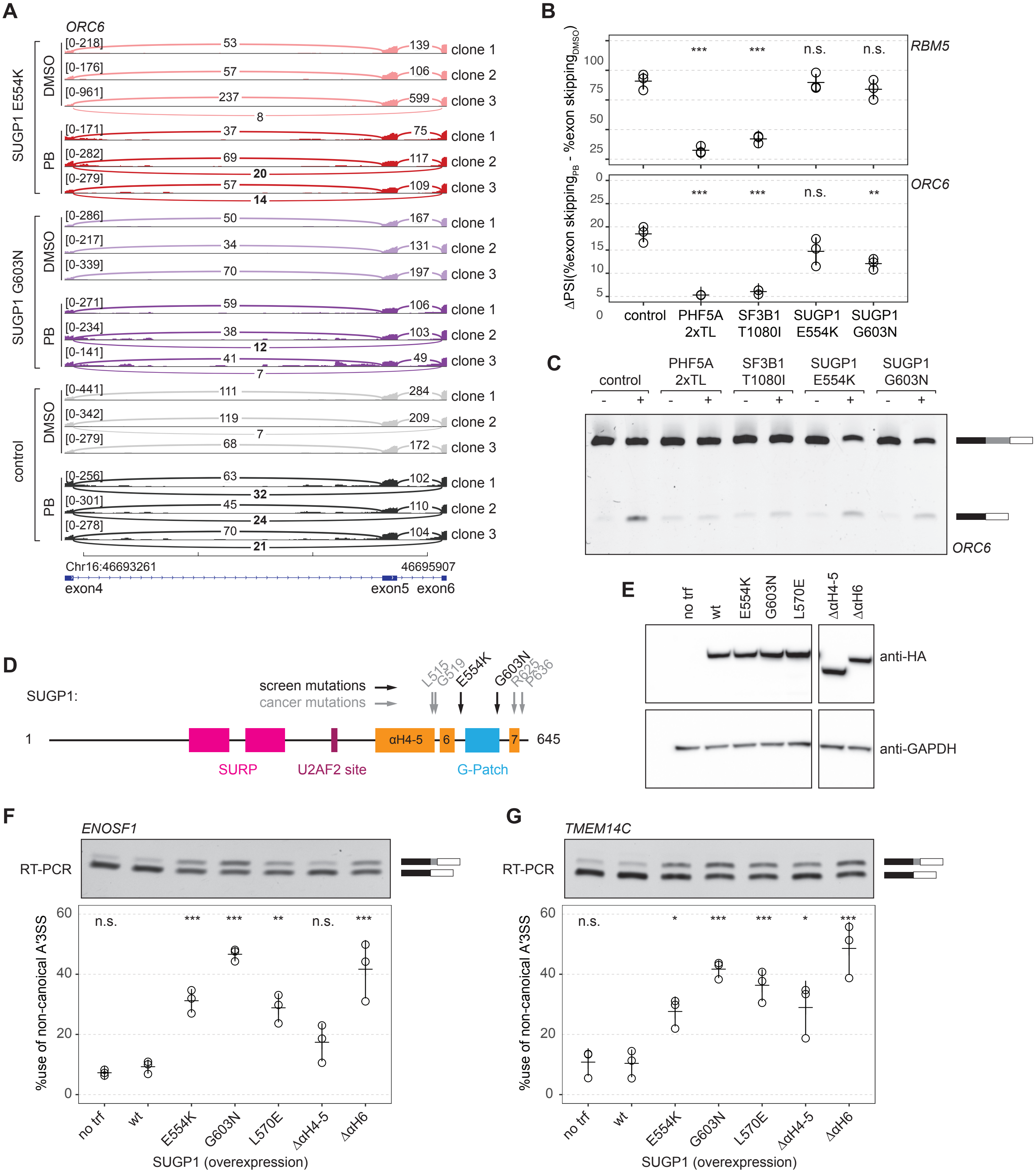

### Supplemental Figure S6

#### Figure S6

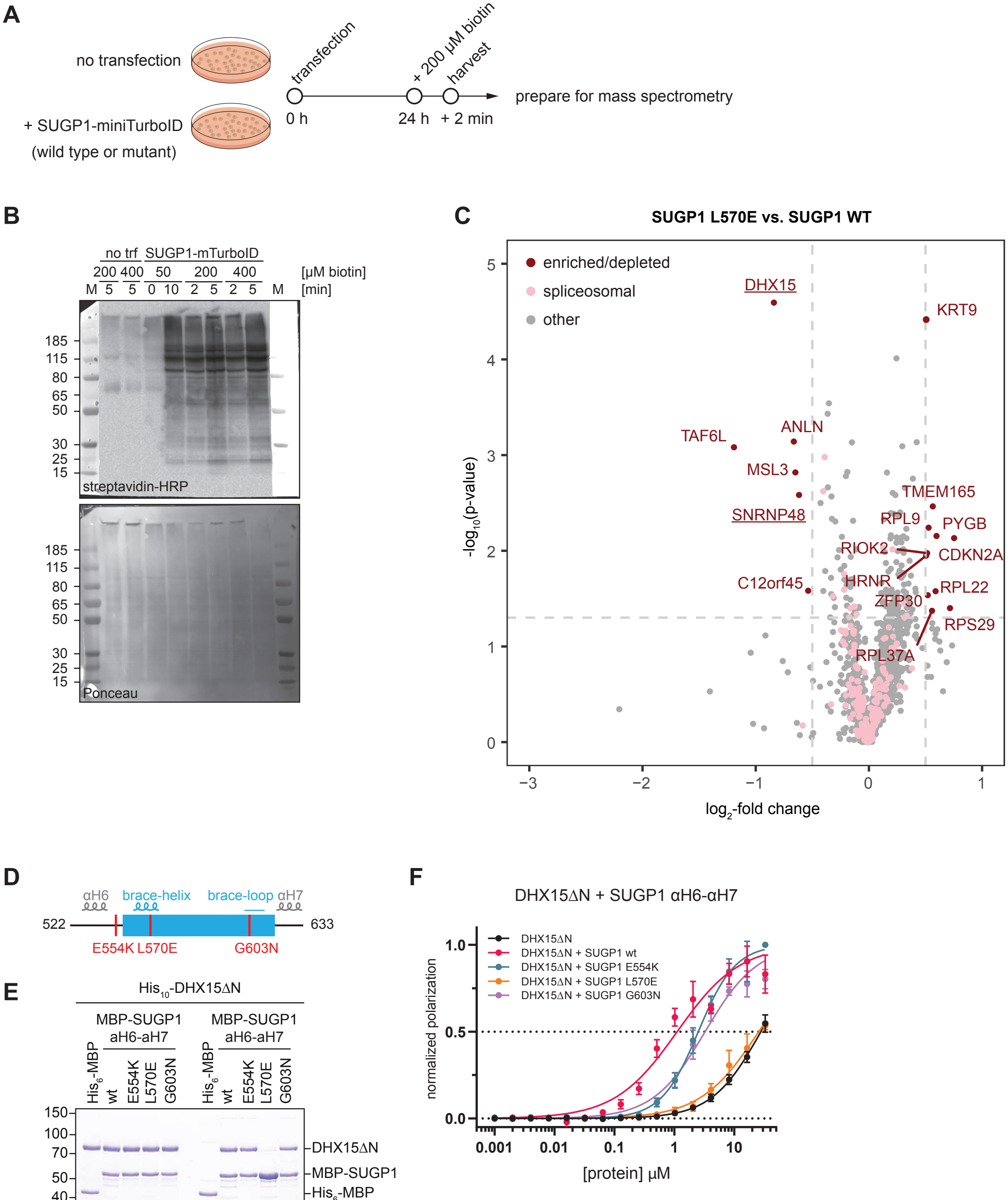


| DHX15ΔN with | $K_d$ [µM] |
|---|---|
| - | 27.9 ± 0.31 |
| SUGP1 αH6-αH7 wt | 1.12 ± 0.16 |
| SUGP1 αH6-αH7 E554K | 2.51 ± 0.06 |
| SUGP1 αH6-αH7 L570E | 25.7 ± 1.21 |
| SUGP1 αH6-αH7 G603N | 3.27 ± 0.30 |
